# EGFR-MAPK adaptor proteins mediate the epithelial response to *Candida albicans* via the cytolytic peptide toxin, candidalysin

**DOI:** 10.1101/2022.03.05.483111

**Authors:** Nicole O. Ponde, Léa Lortal, Antzela Tsavou, Olivia W. Hepworth, Don N. Wickramansinghe, Jemima Ho, Jonathan P. Richardson, David L. Moyes, Sarah L. Gaffen, Julian R. Naglik

## Abstract

*Candida albicans* (*C. albicans*) is a dimorphic human fungal pathogen that can cause severe oropharyngeal candidiasis (OPC, oral thrush) in susceptible hosts. During invasive infection, *C. albicans* hyphae invade oral epithelial cells (OECs) and secrete candidalysin, a pore-forming cytolytic peptide that is required for fungal pathogenesis at mucosal surfaces. Candidalysin induces cell damage and activates multiple MAPK-based innate signaling events that collectively drive the production of downstream inflammatory mediators. The activities of candidalysin are also dependent on the epidermal growth factor receptor (EGFR), but how these signals are integrated is undefined. Here, we identified five essential adaptor proteins as key mediators of the epithelial response to *C. albicans* infection on cultured OECs, including growth factor receptor bound protein 2 (Grb2), Grb2-associated-binding protein 1 (Gab1), Src homology and collagen (Shc), SH2 containing protein tyrosine phosphatase-2 (Shp2) and casitas B-lineage lymphoma (c-Cbl). All these signaling effectors were inducibly phosphorylated in response to *C. albicans*, in a candidalysin-dependent mechanism but additionally required EGFR phosphorylation, matrix metalloproteinases (MMPs) and cellular calcium flux. Of these, Gab1, Grb2 and Shp2 were the dominant drivers of ERK1/2 signaling and production of downstream cytokines. Together, these results identify the key adaptor proteins that drive EGFR signaling mechanisms, which determine oral epithelial responses to *C. albicans*.

## Introduction

Fungal infections are highly under-appreciated contributors of morbidity and mortality in humans, but to date there are no vaccines to any pathogenic fungi (1). *Candida* species are a dominant cause of both superficial and life-threatening fungal infections in humans, with *Candida albicans* being the most prevalent. Though normally a benign member of the oral, vaginal and gut commensal microbiota, disturbances caused by environmental cues, or a compromised immune system result in the overgrowth of *C. albicans*, leading to superficial infections of the mucous membranes, in particularly the oral mucosa, that can significantly lower quality of life and affect millions of individuals annually (2, 3).

An essential feature of *C. albicans* pathogenesis is the formation of hyphae, which cause physical damage to the mucosal epithelium, thereby inducing expression and secretion of innate immune factors such as antimicrobial peptides, alarmins and pro-inflammatory cytokines and chemokines (4). During infection, *C. albicans* hyphae secrete candidalysin, a cytolytic peptide that is critical for pathogenesis at mucosal surfaces (5). Candidalysin is encoded by *ECE1*, which is highly expressed in hyphae, though not required for filamentation. Candidalysin is generated by processing of its parent protein Ece1p by the kexin proteases (6, 7). Candidalysin secretion triggers epithelial cell damage and activation of a mitogen-activated protein kinase (MAPK)-based signaling pathway that leads to secretion of inflammatory mediators and cell survival signals (8–11). The epidermal growth factor receptor (EGFR) is key for mediating the host immune response against several microbial pathogens (12), including *C. albicans* (13–17). However, the downstream signals that orchestrate epithelial responses to this fungus are still incompletely understood.

EGFR, also known as Her1 or ErbB1, belongs to a family of receptor tyrosine kinases (RTKs) that regulate cell survival, proliferation, apoptosis, differentiation and development (18). Dysregulation of EGFR is also associated with tumorigenesis (19, 20). EGFR autophosphorylation is a proximal event that initiates downstream signaling, resulting in recruitment of various adaptors and ultimately activating downstream signaling cascades including MAPK, phosphatidylinositol 3-kinase/Akt/Protein Kinase B (PI3K/Akt), and signal transducers and activators of transcription (STATs). Multiple microbes are known to exploit EGFR during pathogenesis, using adaptor activation and phosphorylation to infiltrate host cells and evade the immune system (12, 21, 22). However, a role for these adaptors in driving immunity to microbes has also emerged (23).

In the present study, we sought to elucidate the role of MAPK signaling and EGFR adaptors in the oral epithelial cell (OEC) response to *C. albicans*. We found that growth factor receptor bound protein 2 (Grb2), Grb2-associated-binding protein 1 (Gab1), Src homology and collagen (Shc), SH2 containing protein tyrosine phosphatase-2 (Shp2, also PTPN11) and casitas B-lineage lymphoma (c-Cbl) are all required to mediate an EGFR-dependent response to *C. albicans* via candidalysin. EGFR kinase inhibitors negatively impact this response, which also requires matrix metalloproteinases and cellular calcium flux. Accordingly, we illuminate the underlying mechanisms of how early EGFR-mediated epithelial responses are orchestrated by the novel fungal toxin candidalysin, with implications for defining the host response to the medically important fungus *C. albicans*.

## Results

### Candida albicans activates EGFR-associated adaptors through secreted candidalysin

In establishment of oropharyngeal candidiasis (OPC, thrush), oral epithelial cells (OECs) respond to the pore-forming toxin candidalysin secreted from *C. albicans* hyphae. Responses require EGFR signals, which lead to the release of antifungal proinflammatory mediators that ultimately coordinate an effective host response (4, 24). However, the underlying mechanisms of the early EGFR molecular events activated in response to *C. albicans* and candidalysin remain incompletely characterized. Accordingly, TR146 buccal epithelial cells were infected with a *C. albicans* (BWP17 + CIp30, parental strain) at a multiplicity of infection (MOI) of 10 for 2 h, a time point known to be optimal for EGFR activation (13). Notably, all proteins known to interact with EGFR were inducibly phosphorylated following infection with *C. albicans* including Gab1 (Y659), Grb2 (p-Tyrosine), Shc (Y317), Shp2 (Y580), c-Cbl (Y774) (Fig. 1A).

**Figure 1.**
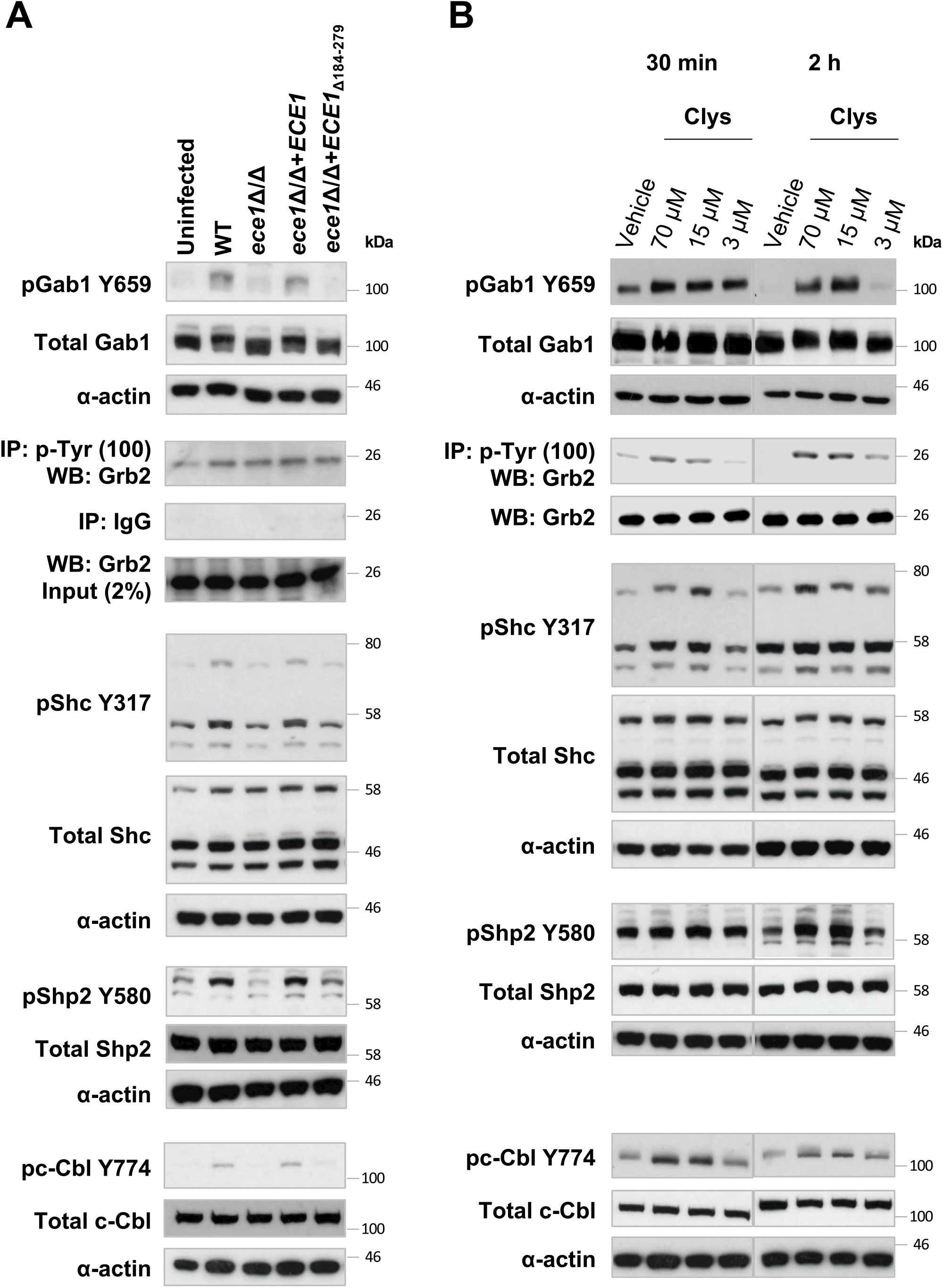
Candidalysin is necessary and sufficient to activate Gab1, Grb2, Shc, Shp2 and c-Cbl adaptors in human oral epithelial cells. (A) Candidalysin activates Gab1, Grb2, Shc, Shp2 and c-Cbl adaptors in human oral epithelial cells (OECs). Immunoblot showing phosphorylation of Gab1, Grb2, Shc, Shp2 and c-Cbl in response to 2 h infection of TR146 OECs with the indicated strains of *C. albicans*. (B) Candidalysin is sufficient to activate Gab1, Grb2, Shc, Shp2 and c-Cbl. Immunoblot showing phosphorylation of Gab1, Grb2, Shc, Shp2 and c-Cbl following stimulation with candidalysin at 30 min and 2 h in oral epithelial cells. Blots are representative of 3 independent experiments.

Since filamentation is required for *C. albicans-*induced epithelial responses (25), we asked whether candidalysin (Clys) was required for adaptor activation. Mutant *C. albicans* strains lacking the candidalysin parent gene *ECE1* (*ece1*Δ/Δ) or with a deletion of the region that encodes candidalysin (*ece1*Δ/Δ+*ECE1*_Δ184–279_) failed to activate Gab1, Shc, Shp2 and c-Cbl (Fig. 1A). However, both mutant strains lacking Clys were able to induce phosphorylation of Grb2, though to a lesser extent than the parental and revertant strains, suggesting candidalysin is not the sole driver of Grb2 phosphorylation. To determine whether Clys alone could activate this panel of adaptors, TR146 cells were stimulated with candidalysin over a time course of 2 h. We treated cells with a strongly lytic concentration (70 μM) that triggers membrane damage, calcium flux and cytokine release, an intermediate sub-lytic concentration (15 μM) that activates epithelial signal transduction through MAPK, p38/MKP-1 and c-Fos and the production of immune regulatory cytokines, and a non-lytic concentration (3 μM) that induces low level secretion of selected cytokines (Moyes et al., 2016). Phosphorylation of Gab1 at Y659 was detectable within 30 min of exposure to all concentrations of candidalysin, with phosphorylation levels increased strongly in response to higher concentrations of candidalysin (70 or 15 μM) at 2 h post treatment (Fig. 1B). Lower concentrations of candidalysin that do not cause cell lysis (3 μM), also induced pGab1 Y659 at 2 h post treatment. In contrast, treatment with 70 or 15 μM candidalysin induced Grb2 phosphorylation within 30 min, which was sustained until 2 h. Increased phosphorylation levels were only observed in cells stimulated with 3 μM candidalysin following 2 h of treatment. Phosphorylation of Shc Y317 was detectable within 30 min following treatment with all concentrations of candidalysin, with a dose-dependent response observed at 30 min and 2 h post stimulation. Together, these data suggest that there is differential temporal organisation of adaptors in response to *C. albicans* and candidalysin, which appears to be an early, coordinated event.

Unexpectedly, the kinetics of Shp2 phosphorylation in response to candidalysin differed from Gab1, Grb2 and Shc, as phosphorylation of Shp2 Y580 was detectable only after 2 h of treatment with 70 or 15 μM candidalysin, with negligible changes in phosphorylation observed with 3 μM (Fig. 1B). These data suggest either that Shp2 may play a role in the response against candidalysin when a significant amount of epithelial damage has been induced, or slower kinetics of Shp2 activity in comparison to the other adaptors. Treatment of TR146 cells with 70 or 15 μM candidalysin triggered strong c-Cbl pY774 activation that was sustained at 30 min and 2 h post stimulation (Fig. 1B). Only very modest changes were observed with 3 μM candidalysin. Collectively, these data show that candidalysin can induce phosphorylation of adaptors in TR146 epithelial cells in a time- and dose-dependent manner, with distinct activation profiles for adaptors in response to lytic versus sub-lytic concentrations of candidalysin.

### Gab1, Grb2, Shc, Shp2 and c-Cbl activation requires EGFR activation, MMPs and calcium flux

While EGFR and the Ephrin family member, EphA2, play a critical role in mediating the innate epithelial response to candidalysin and *C. albicans* (13, 14), it is unknown whether the activation of corresponding adaptors in response to hyphae relies exclusively on EGFR. To determine whether EGFR drives candidalysin-induced adaptor activation, we analyzed the effects of the EGFR-specific kinase inhibitors on the activation of Gab1, Grb2, Shc, Shp2 and c-Cbl. Treatment of TR146 cells with EGFR inhibitors gefitinib or PD153035 reduced *C. albicans*-induced phosphorylation of Gab1, Grb2, Shc, Shp2 and c-Cbl at 2 h post infection (Fig. 2A). Similarly, inhibition of EGFR either abolished or reduced phosphorylation of all adaptors in response to candidalysin (70 or 15 µM), indicating that EGFR activation is indeed required for candidalysin-induced adaptor activity (Fig. 2B).

**Figure 2.**
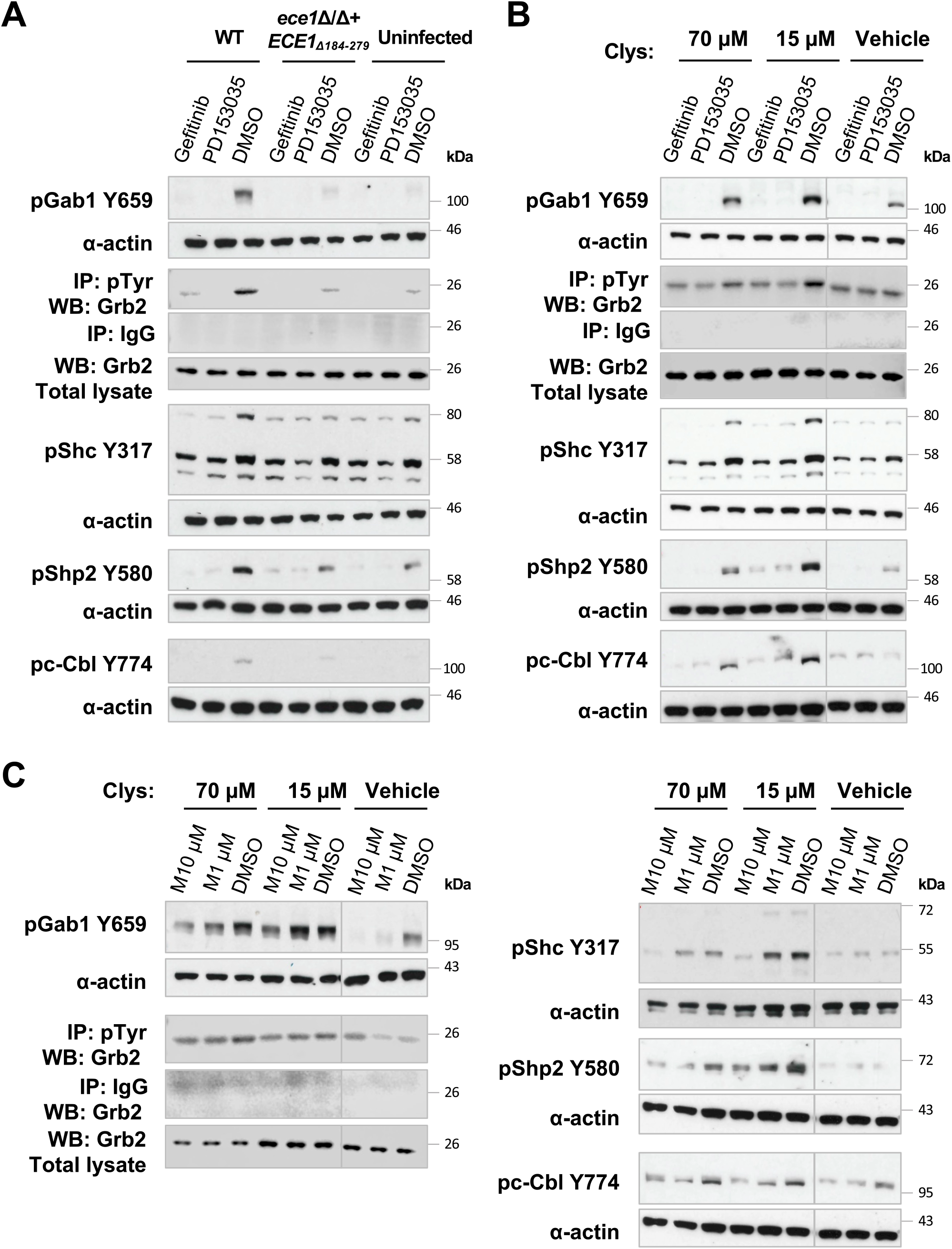
Candidalysin-induced adaptor activation requires EGFR and MMP activity. Inhibition of EGFR activity suppresses (A) *C. albicans-* and (B) candidalysin (Clys)-induced adaptor activation. Use of Gefitinib or PD153035 EGFR TK inhibitors suppresses WT *C. albicans*- and candidalysin-induced activation of pGab1, Grb2, Shc, Shp2 and c-Cbl. (C) MMP inhibition suppresses candidalysin-induced adaptor activity. Pre-treatment of TR146 cells with Marimastat (10 or 1 µM, designated M10 or M1) suppressed candidalysin-induced phosphorylation of Gab1, Grb2, Shc, Shp2 and c-Cbl. Protein lysates were isolated 2 h post stimulation. Blots are representative of 2 (Grb2) or 3 (Gab1, Shc, Shp2 and c-Cbl) independent experiments.

Matrix metalloproteinases (MMPs) are zinc-dependent metallo-endopeptidases that cleave and release EGF-family pro-ligands, therefore activating EGFR signaling (13). Hence, we next assessed the role of MMPs in the activation of adaptors in response to candidalysin. Pre-treatment of TR146 OECs with the pan-MMP inhibitor Marimastat suppressed phosphorylation of Gab1, Grb2, Shc, Shp2 and c-Cbl upon candidalysin stimulation (Fig. 2C) and infection with *C. albicans* (Supp. Fig. S1). Consistent with this, a candidalysin-deficient *C. albicans* strain (*ece1*Δ/Δ+*ECE1*_Δ184–279_), failed to phosphorylate Gab1, Shc and c-Cbl, though induced very modest levels of phosphorylation for Grb2 and Shp2. Additionally, a blocking anti-EGFR monoclonal antibody (cetuximab) reduced/abolished the phosphorylation of Gab1, Grb2, Shc, Shp2 and c-Cbl in response to candidalysin and *C. albicans* infection (Figure 3), demonstrating that candidalysin-induced activation of adaptors is mediated by MMPs, which in turn mediate ligand-induced activation of EGFR. Previously, calcium influx induced by candidalysin activity was also shown to contribute to EGFR phosphorylation (13). Accordingly, pre-treatment of TR146 cells with the calcium chelator, BAPTA-AM also suppressed candidalysin-induced and *C. albicans*-induced EGFR adaptor activation (Supp. Fig. S2). These data show that candidalysin secreted by *C. albicans* activates EGFR-related adaptors and downstream signaling programs in OECs via a mechanism that requires calcium influx, MMP activity and EGFR ligands.

**Figure 3.**
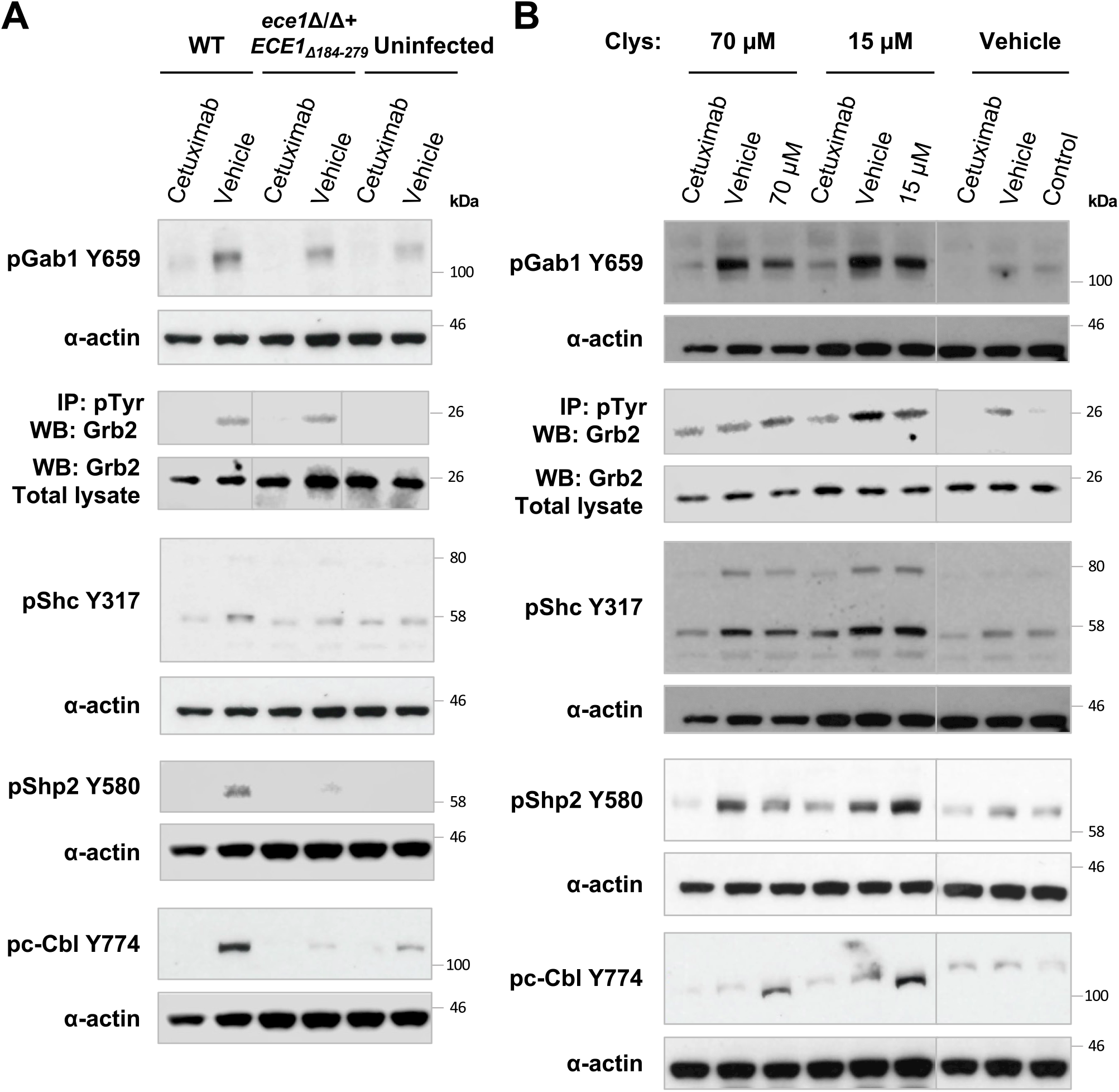
EGFR ligand binding is required for *C. albicans* and candidalysin-induced adaptor activation. Pre-treatment of TR146 cells with the anti-EGFR mAb, Cetuximab suppressed phosphorylation of Gab1, Grb2, Shc, Shp2 and c-Cbl following (A) *C. albicans* infection or (B) candidalysin stimulation. Blots are representative of 2 (Grb2) or 3 (Gab1, Shc, Shp2 and c-Cbl) independent experiments.

### EGFR adaptors mediate candidalysin-induced MAPK signaling responses but are dispensable for cell survival and damage protection

Candidalysin permeabilizes target cell membranes, resulting in damage (release of lactate dehydrogenase (LDH)) and the activation of PI3K/Akt/mTOR and MAPK (MKP1/c-Fos) signaling pathways (5, 26). Notably, the Akt/mTOR pathway is important in mediating cell survival and protection against damage (26) and the MAPK pathway is required for induction of immune responses (5) during epithelial *C. albicans* infection. Furthermore, Gab1, Grb2 and Shp2 adaptors can mediate the cell survival pathway (27).

To determine whether this panel of EGFR adaptors can modulate these cellular responses, TR146 cells were transfected with siRNAs for Gab1, Grb2, Shc, Shp2 and c-Cbl for 48 h, and stimulated with *C. albicans* or candidalysin for 2 h. LDH release from TR146 cells was unaffected by knockdown of all adaptors following treatment with *C. albicans* (Supp. Fig. S3A) or candidalysin (70 or 15 μM) (Supp. Fig. S3B). Accordingly, the candidalysin null strain (*ece1*Δ/Δ+*ECE1*_*Δ184-279*_) caused no detectable change in epithelial integrity. Assessment of the Akt/mTOR pathway similarly showed no change in phosphorylation upon adaptor knockdown (Supp. Fig. S3C). Thus, these EGFR adaptors do not appear to be required to protect epithelial cells from damage or for cell survival in response to *C. albicans* infection and candidalysin. In contrast, siRNA knockdown of Gab1 and Grb2 had a marked impact on c-Fos protein levels and suppressed phosphorylation of MKP1/DUSP1 in response to *C. albicans* and candidalysin (Fig. 4A, 4B). Knockdown of Shc had only a minor impact on c-Fos and MKP1/DUSP1 activation, and the effects of Shp2 or c-Cbl knockdown via siRNA were negligible.

**Figure 4.**
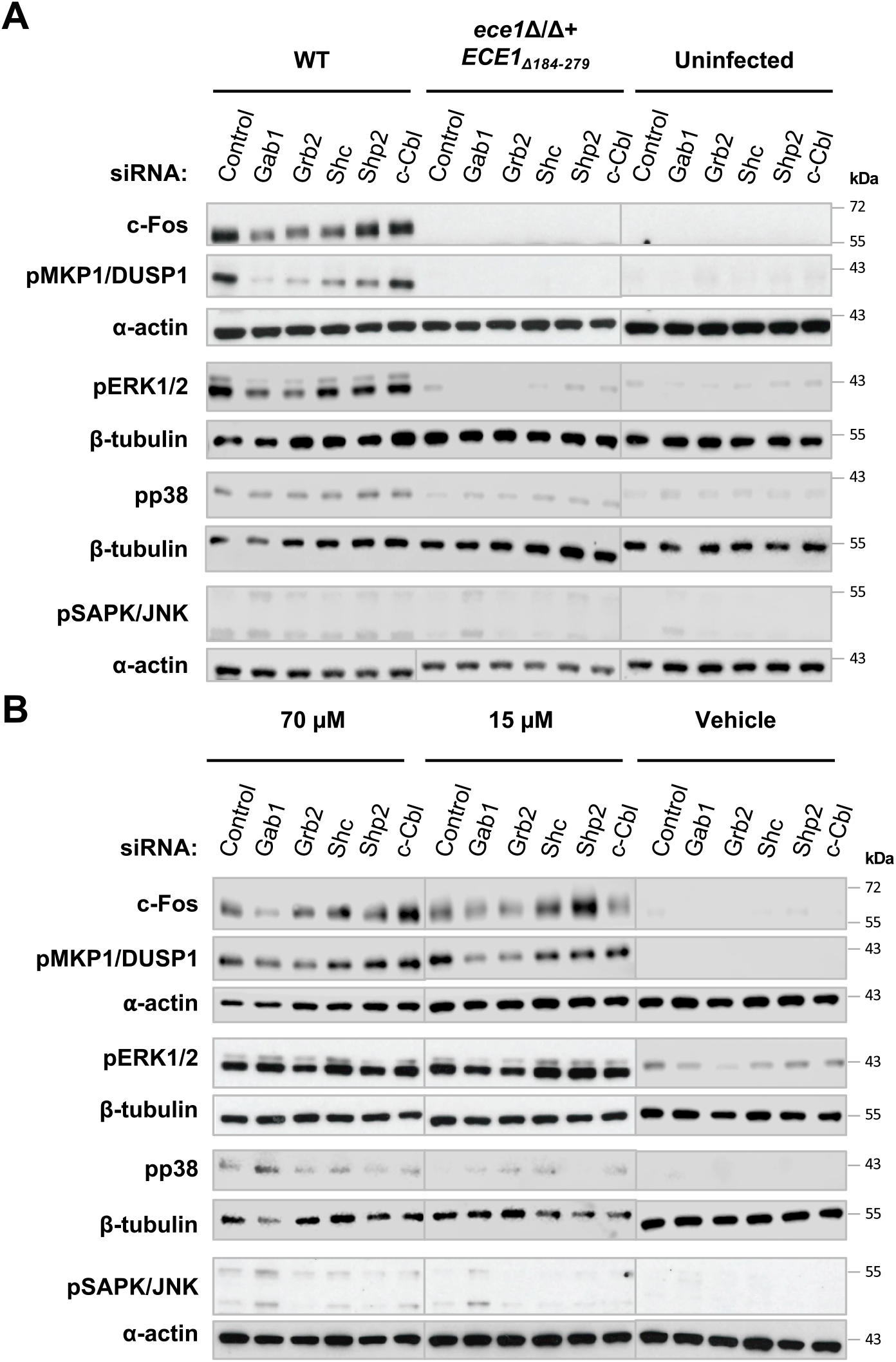
Gab1, Grb2 and Shc adaptors mediate candidalysin-induced MAPK signalling responses. Knockdown of Gab1, Grb2 or Shc suppresses c-Fos expression, MKP1/DUSP1 and ERK1/2 phosphorylation. Following siRNA transfection, cells were infected with the indicated strains (A) or stimulated with candidalysin (B) and lysates were collected after 2 h for western blotting for total c-Fos, phospho-MKP1/DUSP1, phospho-ERK1/2, phospho-p38, phospho-SAPK/JNK or α-actin. Data are representative of 3 independent experiments.

To better understand how Gab1 and Grb2 activate c-Fos and MKP1/DUSP1, we next assessed three major branches of the MAPK signaling cascade, ERK1/2, p38 and SAPK/JNK. In response to *C. albicans*, knockdown of Gab1 and Grb2 had a marked effect on ERK1/2 phosphorylation but not p38 or SAPK/JNK (Fig. 4A, 4B). However, in response to candidalysin, while Grb2 knockdown also showed a decrease in pERK1/2, Gab1 knockdown showed a modest increase in ERK1/2, p38 and SAPK/JNK phosphorylation. As expected, a candidalysin-null strain (*ece1Δ/Δ+ECE1*_*Δ184-279*_) did not trigger MAPK phosphorylation. Thus, Gab1 and Grb2 are required for MAPK activation during *C. albicans* infection, but the Gab1 activation pattern differs when responding to exogenously applied synthetic candidalysin.

Impairing Shp2 via siRNA had no detectable effect on *C. albicans* and candidalysin-induced signals; however, Shp2 knockdown was inefficient (60 %, Supp Fig. S4). Therefore, we addressed this question in an alternate way by treating TR146 cells with a the inhibitor SHP099 HCL, an allosteric inhibitor that selectively and potently targets Shp2. Cells were pre-treated with SHP099 HCL (10 or 5 µM) for 1 h prior to infection and then assessed for cell damage, MAPK and Akt/mTOR pathway activation upon infection. Similar to observations with Shp2 siRNA, LDH release and mTOR phosphorylation from TR146 oral epithelial cells was unaffected by Shp2 inhibition (Supp Fig. S5). However, treatment with SHP099 HCL suppressed *C. albicans*- and candidalysin-induced levels of c-Fos and pMKP1/DUSP1 and pERK1/2, though not pSAPK/JNK or pp38 (Fig. 5). Thus, in addition to a key role for Gab1 and Grb2 adaptors, Shp2 also contributes to ERK1/2 activation in response to *C. albicans* and candidalysin.

**Figure 5.**
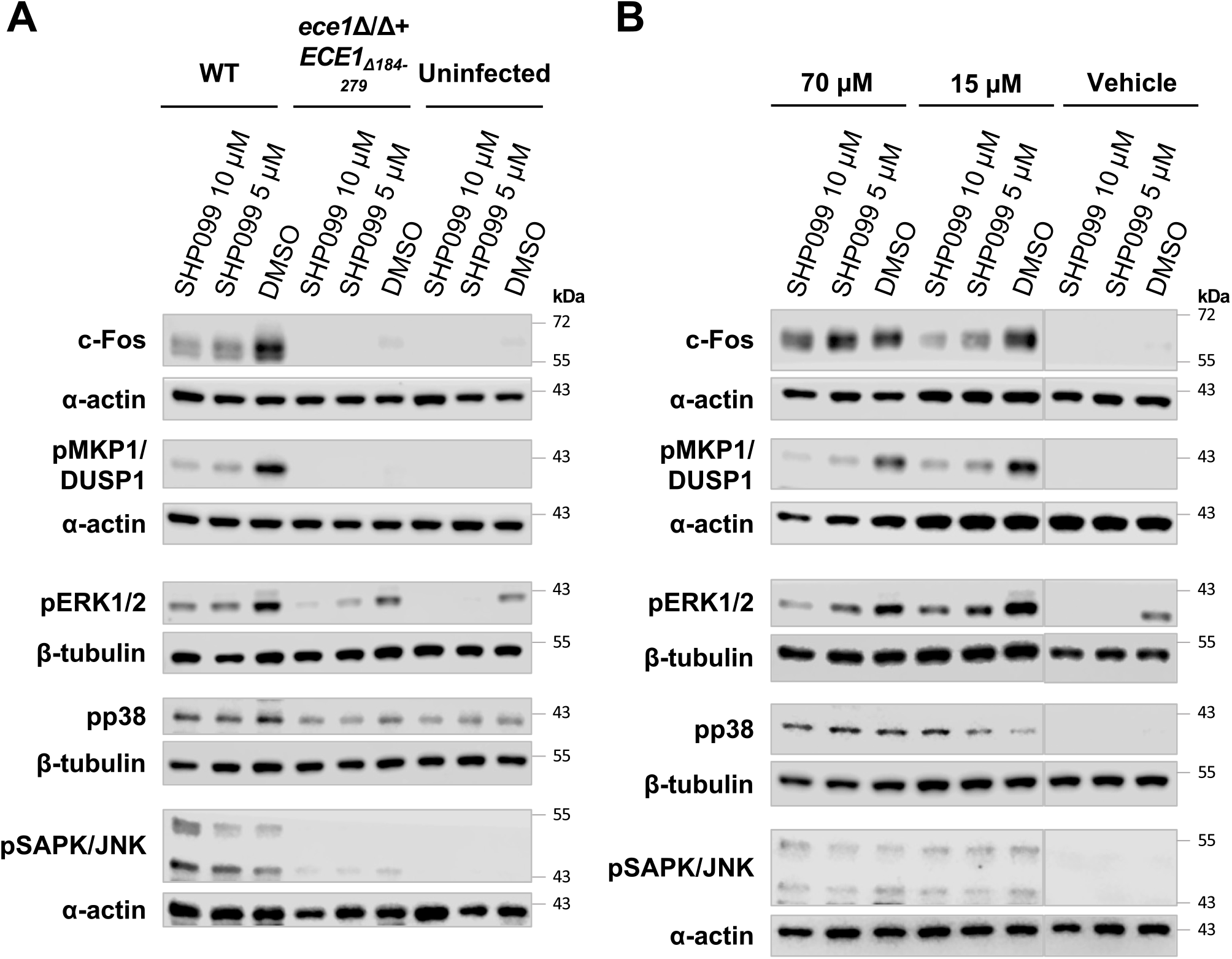
Shp2 mediates candidalysin-induced MAPK signalling responses. Inhibition of Shp2 activity suppresses *C. albicans-* and Candidalysin (Clys)-induced c-Fos, pMKP1/DUSP1 and ERK1/2 activation. Inhibition of Shp2 activity suppresses G-CSF, GM-CSF, IL-1α and IL-1β secretion. TR146 cells were pre-treated with the Shp2 inhibitor, SHP099 HCL for 1 h. Cells were infected with the indicated strains (A) or stimulated with candidalysin (B) and protein lysates isolated at 2 h for western blotting for total c-Fos, phospho-MKP1/DUSP1, phospho-ERK1/2, phospho-p38, phospho-SAPK/JNK or α-actin. Data are representative of 3 independent experiments.

### Gab1, Grb2 and Shp2 are required for production of inflammatory effectors in response to candidalysin

A characteristic end point of candidalysin activity and EGFR activation is the production of inflammatory cytokines and chemokines. Using siRNA knockdown, we evaluated the role of this panel of EGFR adaptors in driving the production of G-CSF, GM-CSF, IL-6, IL-1α and IL-1β from TR146 cells 24 h after *C. albicans* or candidalysin treatment. Knockdown of Gab1, Grb2 or Shc reduced secretion of GM-CSF, G-CSF and IL-1α in response to *C. albicans* (Fig. 6A, 6B, Supp. Fig. 6A) and candidalysin (Supp. Fig. 6B, C, E). Notably, only Gab1 or Grb2 knockdown reduced release of IL-1β and IL-6 in response to *C. albicans* (Fig. 6C, 6D), suggesting that distinct pathways dictate adaptor-mediated cytokine responses during fungal signaling on OECs. Interestingly, knockdown of c-Cbl appeared to increase the release of G-CSF in response to *C. albicans*, and GM-CSF and IL-6 in response to candidalysin (Fig. 6A, Supp. Fig 6C, 6D), possibly suggesting a negative regulatory role for c-Cbl in immune signaling. The Shp2 inhibitor, SHP099 HCL, impaired G-CSF, GM-CSF, IL-1α and IL-β secretion in response to *C. albicans* treatment (Fig. 6E, Supp. Fig. 7A, 7C, 7D). Similarly, Shp2 inhibition significantly reduced G-CSF, GM-CSF and IL-β release in response to candidalysin (Supp Fig. 7E, 7F, 7H). Interestingly, Shp2 inhibition resulted in a significant increase in IL-6 release in response to candidalysin (Fig. 6F).

**Figure 6.**
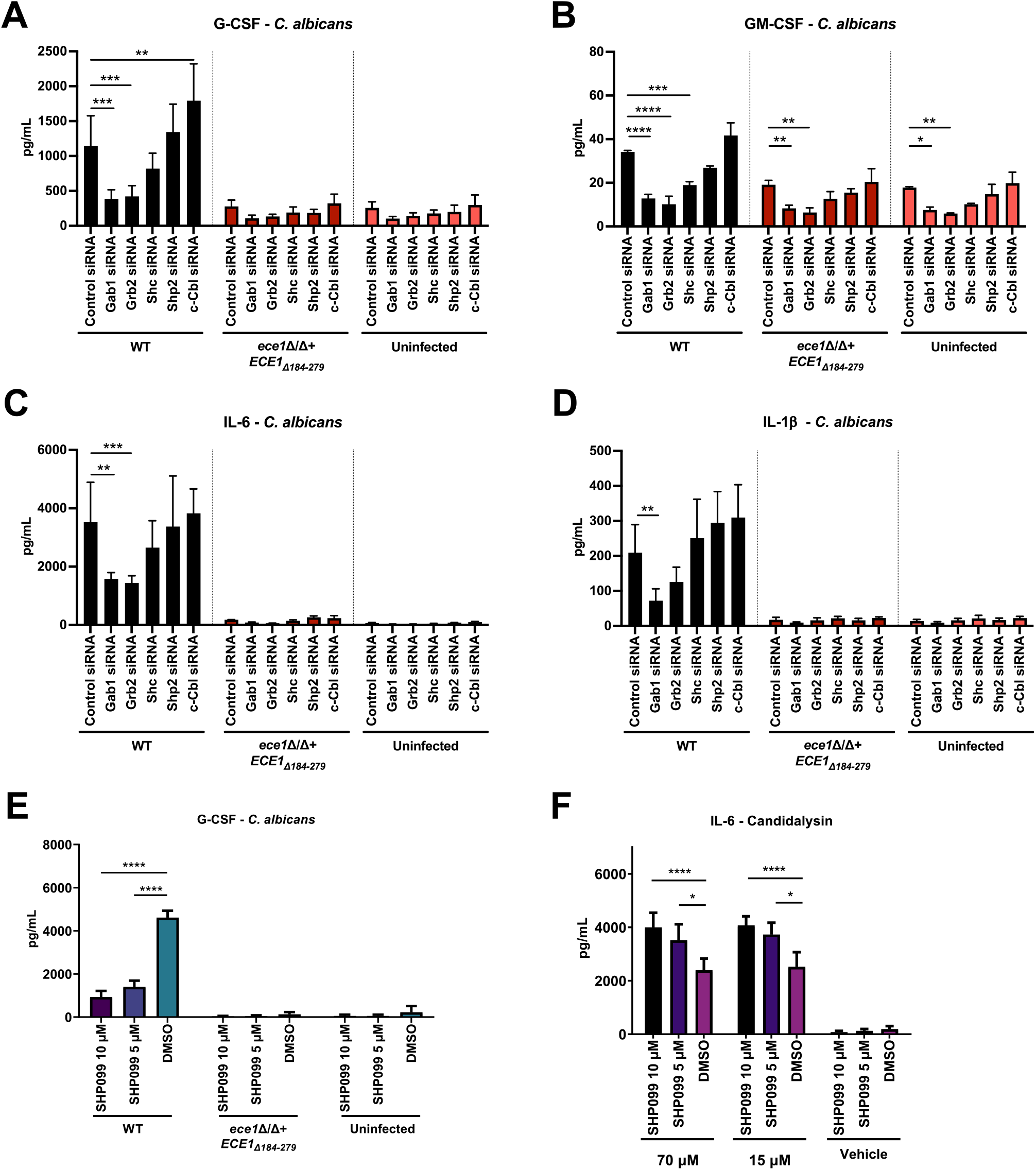
Gab1, Grb2, Shc and Shp2 adaptors mediate *C. albicans- and* candidalysin-induced cytokine secretion in oral epithelial cells. Knockdown of Gab1, Grb2 or Shc or inhibition of Shp2 suppresses G-CSF, GM-CSF, IL-6, and IL-1β secretion. (A-D) Following siRNA knockdown or (E-F) 1 h pre-treatment with the Shp2 inhibitor SHP099HCL, cells were infected with the indicated strains or stimulated with candidalysin and supernatants collected after 24 h. Cytokines were evalluated by Luminex. Data show the mean (± SD) of 3 replicates. Significance was assessed by one way ANOVA with Bonferroni’s multiple comparisons test; *\*P < 0*.*05, **P < 0*.*01, ***P<0*.*001, ****P<0*.*0001*.

OECs respond to candidalysin-induced tissue injury in part by inducing damage associated immune responses, including release of alarmins and antimicrobial peptides (AMPs) such as β-defensins (17). To investigate the role of these adaptors in AMP induction, we incubated TR146 cells with candidalysin following siRNA knockdown of adaptors or inhibition of Shp2 and evaluated *DEFB4A* mRNA (encoding BD2). Adaptor knockdown or inhibition had no significant effect on gene expression in response to candidalysin (Supp. Fig S8A, B). As candidalysin does not stimulate S100 protein release (17), their expression was not assessed. Overall, these data indicate that Gab1, Grb2, Shc and Shp2 adaptors but not c-Cbl play pivotal roles in mediating cytokine but not alarmin innate responses to fungal infection.

## Discussion

The oral epithelium coordinates the early antifungal response to infection through release of cytokines, chemokines and other inflammatory mediators that recruit leukocytes, infiltrating neutrophils and inflammatory monocytes to combat infection. Interactions between epithelial cells and *C. albicans* are critical for host defence mechanisms to mucosal infection. The discovery of the *C. albicans*-derived toxin, candidalysin, was a paradigm shift in understanding how hyphae promote pathogenesis, simultaneously causing target cell damage but also activating innate immune responses that are essential to limit control of this commensal microbe. This discovery led to a myriad of studies revealing how candidalysin activity interfaces with the host immune system and balances commensalism with pathogenicity (6–10, 13, 28–33)

Candidalysin activates the EGFR and MAPK signaling pathways (5, 13), but how these pathways are controlled remained incompletely defined. The EGFR adaptor proteins play a critical role in facilitating signaling between upstream cell surface RTKs, such as EGFR, as well as downstream cellular signaling cascades that are fundamental for a diverse array of cellular processes (34). We show that not only does OEC infection with *C. albicans* activate the Gab1, Grb2, Shc, Shp2 and c-Cbl adaptors, but that this activity is mainly driven by candidalysin. While candidalysin-deficient mutants reduced phosphorylation of Grb2 in OECs compared to candidalysin-producing strains, there were still residual levels of phosphorylation remaining, suggesting that candidalysin is not the only driver of Grb2 activity during infection. This may be due to interactions between fungal proteins on the surface of invading hyphae, and host epithelial cell receptors, EphA2, EGFR and globular C1q receptor (gC1qR) which are known to mediate proinflammatory responses to *C. albicans* (14, 35, 36).

The distinct temporal activity of these EGFR adaptors suggest they may mediate the early, coordinated response of OECs to ensure appropriate signaling during fungal infection. Notably, no studies have implicated EGFR adaptor activation in response to fungal infection and very few in bacterial infections (21, 22, 37). In addition to EGFR, Gab1, Grb2, Shc, Shp2 and c-Cbl are essential for signaling responses of RTKs, including EphA2, c-Met and other Her-family members (38–40). The near abolishment of adaptor activation in response to EGFR inhibition confirmed EGFR is a critical receptor for adaptor protein activation and downstream responses to *C. albicans* via candidalysin. Previously, we reported that inhibition of EGFR reduces MAPK signaling and subsequent immune responses during infection (13). Interestingly, our results show that some, but not all, adaptors orchestrate the downstream mechanisms of epithelial immunity to infection. Recent work has demonstrated that Grb2 is associated with EGFR in response to *C. albicans* infection (36), and sustained activation of both EGFR and EphA2 by candidalysin and the hyphal adhesin, Als3, is required for proinflammatory responses during *C. albicans* infection (14). However, unlike candidalysin, recombinant Als3 does not induce cell damage, activate c-Fos/MKP1/DUSP1, or promte cytokine responses (41), which are downstream of EGFR signaling, suggesting Als3 does not directly activate these EGFR adaptors. Additionally, it should be noted that an *ALS3*-deficient strain is poor at forming the invasion pocket, which is required to enable efficient delivery and activity of candidalysin (8). Together, these data indicate that Als3 does not directly activate EGFR adaptors, and that its role in driving epithelial signaling and/or immune responses is probably due to its role in facilitating the formation of the invasion pocket to permit efficient secretion of candidalysin, which activates these processes.

Candidalysin induces EGFR activation by activating matrix metalloproteinases (MMPs), which cleave EGFR pro-ligands that subsequently bind EGFR, and inducing calcium influx (5, 13). Various MMPs activate EGFR in response to a diverse array of stimuli (42, 43). Additionally, MMP activation and subsequent cleavage and release of EGFR ligands are involved in the activation of EGFR in response to *Helicobacter pylori* and *Clostridium difficile* (44, 45). The role of MMPs in infectious diseases, such as mycobacterial infection and *Paracoccidioides brasiliensis* infection has also been reported (46, 47). We now demonstrate that MMPs mediate antifungal immune responses via EGFR through the adaptors described herein.

Candidalysin is the first pore-forming toxin (PFT) identified in a human fungal pathogen. Like analogous toxins in other organisms, candidalysin causes a rapid influx of intracellular calcium, thought to be a pivotal step in the activation of downstream signaling events (48). Calcium influx triggered by bacterial PFTs is also a key factor in the induction of cellular responses that limit infection and promote survival (49, 50). We found that chelation of calcium reduced the activation of adaptors in response to *C. albicans* and candidalysin. As such, our results demonstrate a role for calcium flux induced by candidalysin in inducing EGFR adaptor activation and epithelial immune responses during *C. albicans* infection.

Candidalysin induces cell stress in OECs, with the release of alarmins, ATP and antimicrobial peptides (AMPs) (17). Our findings reveal that these adaptors play a negligible role in expression of a key host-protective AMP, *DEFB4A* (human BD-2). Interestingly, Shc activation has been associated with cellular stress responses (51), and Shp2 is critical in regulating the stress response to aryl hydrocarbon receptor activation and calcium dynamics in mast cells (52). Though this aspect of adaptor activation via cellular stress was not assessed here, stress-induced calcium flux is a key biological response to bacterial PFTs, and it is likely that candidalysin induces stress-associated signals in OECs during *C. albicans* infection, which may activate EGFR adaptors to mediate this specific response to candidalysin.

This work establishes roles for multiple EGFR-related adaptors in the response to candidalysin secreted from *C. albicans*. Notably, four of these proteins, Gab1, Grb2, Shc and Shp2, were required for G-CSF and GM-CSF secretion (Fig. 6A, 6B, S7A, S7E, S7F). The secretion of IL-1α was reduced when Gab1 or Grb2 were knocked down via siRNA (Fig. S6A) or when the activity of Shp2 was inhibited chemically (Fig. S7C). In a similar manner, IL-1β and IL-6 secretion were reduced upon siRNA knockdown of Gab1 but not Grb2 in response to infection or candidalysin treatment (Fig. 6D). Inhibition of Shp2 effectively reduced IL-1β but not IL-6. These distinct roles for each adaptor in mediating the secretion of IL-1α, IL-1β and IL6 indicates a potentially extensive and complex network of regulation of these cytokines in response to infection (Fig. 7) (Table 1).

**Figure 7.**
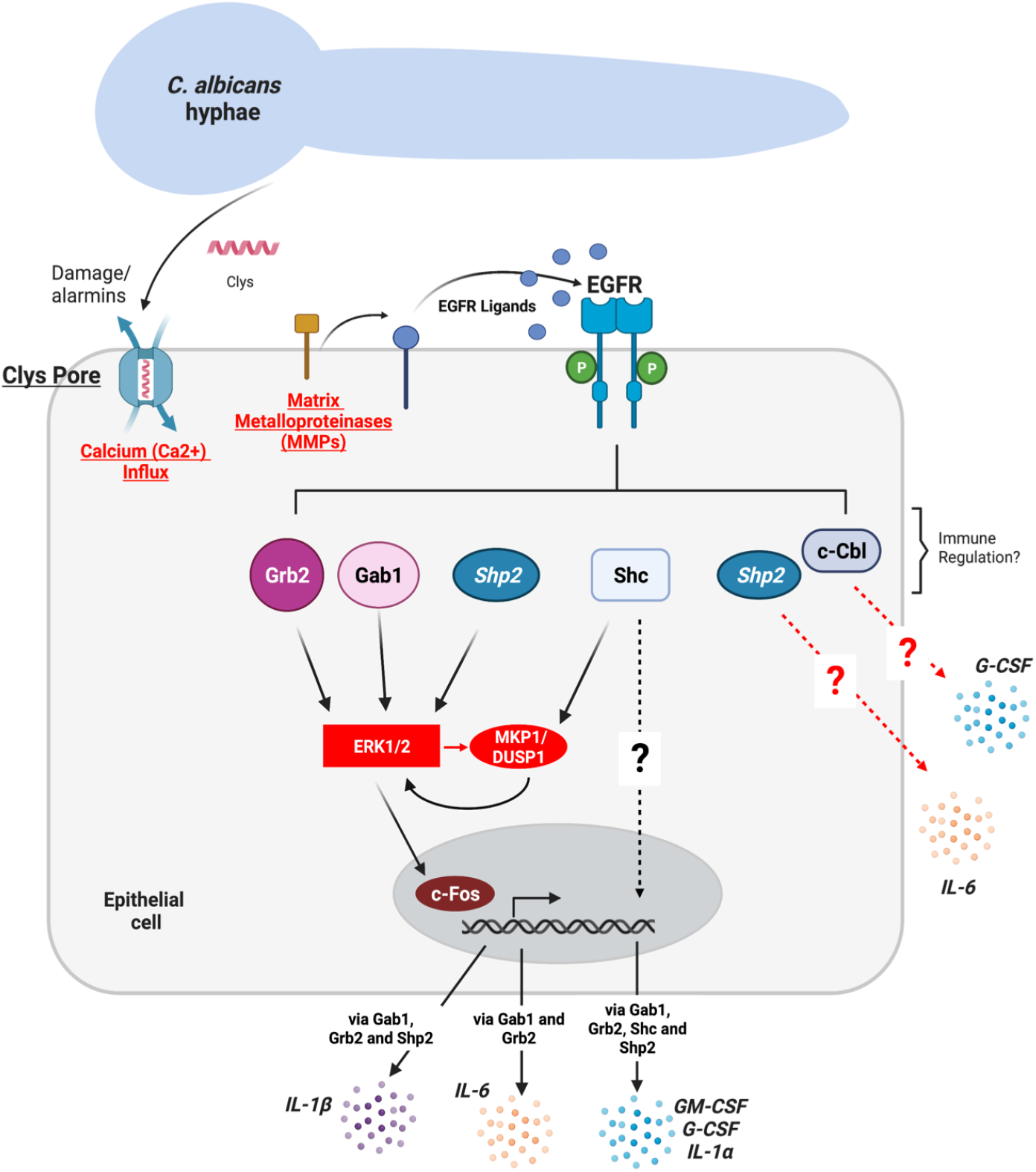
Gab1, Grb2, Shc and Shp2 adaptors mediate candidalysin-induced signaling events that trigger oral epithelial cytokine responses. *C. albicans* infections are initiated by increased fungal burden, associated hypha formation and secretion of the pore-forming toxin candidalysin (Clys). When accumulated at high concentrations, candidalysin interacts with an epithelial cell membrane to form pore-like structures that result in membrane damage, alarmin release and calcium influx. These events lead to activation of matrix metalloproteinases (MMPs) and the cleavage/release of epidermal growth factor receptor (EGFR) ligands. EGFR activation leads to the recruitment and activation of the Gab1, Grb2, Shc, Shp2 and c-Cbl adaptor proteins which in turn mediate MKP1/DUSP1 and c-Fos activation via the MAPK-ERK1/2 signalling pathway. Cumulatively, this leads to induction of GM-CSF and G-CSF (promoting neutrophil recruitment), as well as IL-6, IL-1α and IL-1β which mediate the subsequent stages of the innate and adaptive immune responses. Figure created with Biorender.com

**Table 1.** Summary of EGFR-MAPK adaptor knockdown and inhibition phenotypes.

| Adaptor Knocked Down/Inhibited |  |  |  |  |  |
| --- | --- | --- | --- | --- | --- |
| Cytokine | Gab1 | Grb2 | Shc | Shp2 | c-Cbl |
| G-CSF | ↓↓ | ↓↓ | ↓↓ | ↓↓ | ↑↑ |
| GM-CSF | ↓↓ | ↓↓ | ↓↓ | ↓↓ | - |
| IL-6 | ↓↓ | ↓↓ | - | ↑↑ | - |
| IL-1α | ↓↓ | ↓↓ | ↓ | ↓ | ↓ |
| IL-1β | ↓↓ | ↓ | - | ↓ | - |

↓↓ Significant Reduction ↓ Reduced - No change ↑ Increased ↑↑ Significant Increase

Few studies show that Gab1, Grb2 or Shp2 mediate such specific cytokine responses. However, activation of Gab1 and Shp2 can transmit signals to the ERK1/2 signaling cascade to activate the cytokine receptor, gp130 (53), and activation of Shc, Grb2 and Shp2 mediate MAPK activation via the G-CSF receptor (54). Both G-CSF and GM-CSF are strong neutrophil recruitment cytokines required for the resolution of *C. albicans* infections (55, 56). The decrease in ERK1/2 phosphorylation in combination with G-CSF and GM-CSF reduction upon Gab1 and Grb2 knockdown suggests that both adaptor proteins control the OEC immune response to candidalysin, which may subsequently mediate neutrophil recruitment during *C. albicans* infection *in vivo*.

EGFR adaptor activation and the OEC response to fungal infection are not well understood. Our data show that ERK1/2 is the major pathway activated by the EGFR adaptors, rather than p38 or SAPK/JNK, which is in agreement with work showing that candidalysin activates p38 independently of EGFR and ERK1/2 and that the EGFR-ERK1/2-c-Fos pathway is downstream of p38 (11). During infection with *C. albicans*, MAPK signaling is regulated by MKP1/DUSP1, with ERK1/2 mediating a negative feedback loop to coordinate the innate immune response (25). These data suggest a dual role for ERK1/2 signaling in response to *C. albicans* and candidalysin; (i) as a mediator of epithelial immune responses and (ii) a regulator of MAPK signaling responses. Interestingly, we only observed a partial reduction in c-Fos total protein when Gab1, Grb2, Shc or Shp2 were knocked down or inhibited during fungal exposure, suggesting that additional unidentified factors can promote cytokine responses against *C. albicans* and candidalysin.

Although Gab1, Grb2 and Shc mediate cytokine responses in response to *C. albicans* and candidalysin, knockdown of c-Cbl showed either no effect or an increase in G-CSF, GM-CSF and IL-1β release. Additionally, inhibition of Shp2 resulted in an increase in IL-6 secretion, most notable with candidalysin. This suggests a role for c-Cbl and Shp2 as possible negative regulators of EGFR signaling and immune induction. Shp2 has been shown to promote bacterial clearance following post-influenza *Staphylococcus aureus* pneumonia (57) and orchestrate macrophage function against pulmonary bacterial infection (58), but its role in fungal infections is largely undefined. However, Shp2 is known to negatively regulate IL-6 release in macrophages in response to TLR ligands (59), thus, we anticipate that Shp2 has a similar function during OEC infections. The c-Cbl adaptor is important in EGFR trafficking and regulation (60, 61); notably, ubiquitination of EGFR by c-Cbl is implicated in clathrin-independent endocytosis of EGFR and downstream signaling (62). Additionally, though the p38 MAPK pathway is pivotal in EGFR endocytosis (63), little is known about the role of p38 in EGFR cellular localization in response to infection. It is likely that the complex dynamics of EGFR trafficking during fungal infection impacts EGFR signaling and consequently the OEC immune response.

In summary, these data indicate that the EGFR-associated adaptors Gab1, Grb2, Shc, Shp2 and c-Cbl play distinct roles in regulating OEC responses to *C. albicans* infections (Fig. 7). Though these adaptors are key mediators supporting microbial infection, relatively few studies evaluate these adaptors as regulators of the epithelial immune response. As such, it is likely that EGFR-associated adaptors act as sentinel regulators of epithelial immune responses during mucosal infections.

## Experimental Procedures

### Candida albicans strains and Candidalysin

*C. albicans* strains were previously described (5) and maintained on yeast extract peptone dextrose (YPD) agar and cultured in YPD at 30°C. Fungal cells were harvested by centrifugation, washed twice with PBS and diluted in appropriate media. Candidalysin (SIIGIIMGILGNIPQVIQIIMSIVKAFKGNK; also termed Ece1-III_62-92K_) was synthesized by Peptide Protein Research Ltd, UK and stored frozen at 3.021 mM (10 mg/mL) in sterile water.

### Cell Culture

TR146 human buccal epithelial squamous cell carcinoma cells (64) were from European Collection of Authenticated Cell Cultures (ECACC) and cultured in Dulbecco’s Modified Eagle Medium Nutrient Mixture F-12 HAM (DMEM/F-12)(Gibco) supplemented with 15% (Gibco) and 1% (v/v) penicillin-streptomycin)(Sigma-Aldrich). Serum-free DMEM/F-12 replaced normal growth medium 24 h before and during cell stimulations.

### Reagents

Inhibitors were from Santa Cruz Biotechnology (Gefitinib), Selleckchem (PD153035, SHP099HCL) and Tocris Bioscience (Marimastat). BAPTA-AM was from Selleckchem. Inhibitors were reconstituted in DMSO and stored frozen. Cetuximab was a gift from Guy’s Hospital Cancer Centre and stored at 4°C. TR146 cells were incubated with inhibitors for 1 h prior to *C. albicans* infection, candidalysin treatment. Antibodies sources: Cell Signaling Technology: pGab1 Y659 (#12745S), Gab1 (#3232S), Grb2 (#3972S), pShc Y317 (#2431S), Shc (#2432S), pShp2 Y580 (#3703S), Shp2 (#3752S), pc-Cbl Y774 (#3555S), c-Cbl (#8447S), pMKP1/DUSP1 S359 (#2857S), c-Fos (#2250S), pp44/42 MAPK (ERK1/2) T202/Y204 (#4370S), pp38 MAPK T180/Y182 (#4511S), pSAPK/JNK T183/Y185 (#4668S), pAkt S473 (#4060S), pMTOR S2448 (#5536S) and β-Tubulin (#2146S); Millipore: mouse α-actin (#MAB1501); Jackson ImmunoResearch: goat anti-mouse IgG (#115-035-062), goat anti-rabbit IgG (#111-035-144) horseradish peroxidase (HRP).

### RNA Silencing

RNAi primers were from ThermoFisher Scientific. TR146 cells were transfected with 5 or 10 nM of Gab1 (s5463), Grb2 (s226232), Shc (s12813), Shp2 (s11525), c-Cbl (s2476) (Table S1) or a negative control siRNA (Silencer™ Negative Control No. 2 siRNA) using HiPerfect Transfection Reagent (Qiagen) per manufacturers’ instructions. The extent of knockdown was verified after 48 h by immunoblotting or qPCR.

### qPCR

RNA was isolated with RNeasy Mini Kits (Qiagen). Complementary DNA was synthesized using iScript cDNA synthesis Kit (Bio-Rad). Real-time qPCR was performed with SsoAdvanced Universal SYBR Green Supermix (Bio-Rad) on a CFX Opus 96 Real-Time PCR Instrument (Bio-Rad), normalized to *GAPDH*. Primers were from QuantiTect Primer Assays (Qiagen).

### Immunoblotting

TR146 cells were incubated with the indicated fungal strains or treated with candidalysin (70, 15 or 3 µM) for the indicated timepoints. Cells were rinsed with PBS and lysed with RIPA buffer (50 mM Tris-HCl pH 7.5, 150 mM NaCl, 1% (v/v) Triton X-100, 1% (w/v) sodium deoxycholate, 0.1% (w/v) SDS, 20 mM EDTA) containing protease and phosphatase inhibitors (Sigma-Aldrich). Cells were detached with a scraper and lysates incubated on ice for 30 min. 5-10 µg of whole protein extracts were separated on Bolt™ 4 to 12%, Bis-Tris precast gels and transferred to nitrocellulose (Bio-Rad). Membranes were probed for target proteins with specific antibodies against phosphorylated or total proteins and secondary antibodies before being developed using Immobilon chemiluminescent substrate (Millipore) and visualized using the Odyssey Infrared Imaging System (LI-COR).

### Immunoprecipitation

TR146 cells were serum starved and infected with *C. albicans* strains at an MOI of 10 or treated with candidalysin (70, 15 or 3 µM) for the indicated times. For inhibitor experiments, cells were pre-treated with inhibitors 1 h prior to infection or stimulation. Cells were washed with PBS and lysed with buffer (25 mM Tris-HCl pH 7.4, 150 mM NaCl, 1% Nonidet P-40 (NP-40), 1 mM EDTA, 5% glycerol) supplemented with protease and phosphatase inhibitors (Sigma-Aldrich) for 30 min. Lysates were centrifuged at 13,000 x g at 4°C for 10 min. Approximately 100 µL of total lysate was used as loading controls and the remaining lysates precleared with 25 µL of protein A/G magnetic beads (Pierce, Thermo Fisher). A mouse phosphotyrosine antibody conjugated to a magnetic bead (#8095S, Cell Signaling Technology) or control IgG antibody (#5873, Cell Signaling Technology) was added to 250 μg of protein and incubated with rotation overnight at 4°C. Samples were pelleted using a magnetic rack, washed four times with ice cold lysis buffer and protein eluted with 20 μL of 3x SDS buffer containing DTT by heating at 95°C for 5 min. Eluted proteins were separated by SDS-PAGE, and analyzed as described above.

### Quantification of cell damage (LDH assay)

A Cytox 96 Non-Radioactive Cytotoxicity Assay Kit (Promega) was used to measure the activity of lactate dehydrogenase (LDH) in cell culture supernatants as described (13, 17). TR146 cells were seeded and stimulated with *C. albicans* or candidalysin. Following each incubation time, exhausted culture medium was collected and analysed for LDH activity as an indicator of *C. albicans* or candidalysin-induced membrane disruption. Assay was conducted according to the manufacturer’s instructions and recombinant L-Lactic Dehydrogenase from porcine heart (Sigma-Aldrich) was used to generate a standard curve. Samples were measured for LDH following collection from culture plates.

### Cytokine secretion

Cytokine levels in cell culture supernatants were determined as previously described (13). Following infection with *C. albicans* at an MOI of 0.01 or stimulation with candidalysin, conditioned culture medium was collected after 24 h and assessed by Luminex assay (Bio-techne) and a Bioplex-200 machine (Bio-Rad). Bioplex manager 6.1 software was used to determine analyte concentrations.

### Statistics

Data were compared by ANOVA with Bonferroni’s multiple comparisons test using GraphPad Prism (v. 8) software. *P* values < 0.05 were considered significant.

## Data Availability

All data on adaptor phosphorylation, inhibition and knockdown assays and cytokine expression are contained within this manuscript.

## Supporting Information

This article contains supporting information.

## Acknowledgments

We thank Bernhard Hube for providing fungal strains and members of the Centre for Host-Microbiome Interactions at King’s College London and the Division of Rheumatology and Clinical Immunology at the University of Pittsburgh for helpful suggestions.

## Author contributions

N.O.P, L.L, A.T, O.W.H., D.N.W., investigation; N.O.P., methodology; N.O.P., writing-original draft; S.L.G, J.P.R. and J.N., writing-review and editing; J.P.R., D.L.M. and J.R.N. supervision; D.L.M, J.H, S.L.G, J.R.N., conceptualization; S.L.G and J.R.N., funding acquisition; J.R.N., project administration.

## Funding and additional information

S.L.G was supported by the National Institutes of Health (DE022550). J.R.N. was supported by the Wellcome Trust (214229_Z_18_Z), National Institutes of Health (DE022550), and the National Institute for Health and Care Research (NIHR) Biomedical Research Centre based at Guy’s and St Thomas’ NHS Foundation Trust and King’s College London and/or the NIHR Clinical Research Facility (IS-BRC-1215-20006). D.L.M was supported by the National Institutes of Health (DE022550), and the UKRI-BBSRC (BB/S016899/1). The views expressed are those of the author(s) and not necessarily those of the NHS, the NIHR or the Department of Health and Social Care.

## ^1^ Abbreviations

AKT/PKB: Protein Kinase B
c-CBL: Casitas B-Lineage Lymphoma
ECE1: Extent of Cell Elongation
ERK: Extracellular Signal-Regulated Kinase
EGFR: Epidermal Growth Factor Receptor
GAB1: Grb2-associated-binding Protein 1
GRB2: Growth Factor Receptor-Bound Protein 2
MAPK: Mitogen-Activated Protein Kinase
MMPs: Matrix Metalloproteinases
OEC: oral epithelial cells
OPC: Oropharyngeal Candidiasis
SAPK/JNK: Stress-activated protein kinases/Jun amino-terminal kinases
SHC: Src Homology 2 And Collagen Protein
SHP2: SH2 Containing Protein Tyrosine Phosphatase-2

## References

1. Brown, G. D., Denning, D. W., Gow, N. A. R., Levitz, S. M., Netea, M. G., and White, T. C. (2012) Hidden killers: Human fungal infections. Science Translational Medicine. 4, 165rv13–165rv13

2. Romani, L. (2011) Immunity to fungal infections. Nat Rev Immunol. 11, 275–288

3. Verma, A., Gaffen, S. L., and Swidergall, M. (2017) Innate Immunity to Mucosal Candida Infections. J Fungi (Basel). 10.3390/JOF3040060

4. Pellon, A., Sadeghi Nasab, S. D., and Moyes, D. L. (2020) New Insights in Candida albicans Innate Immunity at the Mucosa: Toxins, Epithelium, Metabolism, and Beyond. Frontiers in Cellular and Infection Microbiology. 10.3389/fcimb.2020.00081

5. Moyes, D. L., Wilson, D., Richardson, J. P., Mogavero, S., Tang, S. X., Wernecke, J., Höfs, S., Gratacap, R. L., Robbins, J., Runglall, M., Murciano, C., Blagojevic, M., Thavaraj, S., Förster, T. M., Hebecker, B., Kasper, L., Vizcay, G., Iancu, S. I., Kichik, N., Häder, A., Kurzai, O., Luo, T., Krüger, T., Kniemeyer, O., Cota, E., Bader, O., Wheeler, R. T., Gutsmann, T., Hube, B., and Naglik, J. R. (2016) Candidalysin is a fungal peptide toxin critical for mucosal infection. Nature. 532, 64–68

6. Liu, J., Willems, H. M. E., Sansevere, E. A., Allert, S., Barker, K. S., Lowes, D. J., Dixson, C., Xu, Z., Miao, J., DeJarnette, C., Tournu, H., Palmer, G. E., Richardson, J. P., Barrera, F. N., Hube, B., Naglik, J. R., and Peters, B. M. (2021) A variant ECE1 allele contributes to reduced pathogenicity of Candida albicans during vulvovaginal candidiasis. PLoS Pathogens. 10.1371/JOURNAL.PPAT.1009884

7. Richardson, J. P., Mogavero, S., Moyes, D. L., Blagojevic, M., Krüger, T., Verma, A. H., Coleman, B. M., de La Cruz Diaz, J., Schulz, D., Ponde, N. O., Carrano, G., Kniemeyer, O., Wilson, D., Bader, O., Enoiu, S. I., Ho, J., Kichik, N., Gaffen, S. L., Hube, B., and Naglik, J. R. (2018) Processing of Candida albicans ece1p is critical for Candidalysin maturation and fungal virulence. mBio. 10.1128/mBio.02178-17

8. Mogavero, S., Sauer, F. M., Brunke, S., Allert, S., Schulz, D., Wisgott, S., Jablonowski, N., Elshafee, O., Krüger, T., Kniemeyer, O., Brakhage, A. A., Naglik, J. R., Dolk, E., and Hube, B. (2021) Candidalysin delivery to the invasion pocket is critical for host epithelial damage induced by Candida albicans. Cellular Microbiology. 23, e13378

9. Allert, S., Förster, T. M., Svensson, C. M., Richardson, J. P., Pawlik, T., Hebecker, B., Rudolphi, S., Juraschitz, M., Schaller, M., Blagojevic, M., Morschhäuser, J., Figge, M. T., Jacobsen, I. D., Naglik, J. R., Kasper, L., Mogavero, S., and Hube, B. (2018) Candida albicans-Induced Epithelial Damage Mediates Translocation through Intestinal Barriers. mBio. 10.1128/MBIO.00915-18

10. Blagojevic, M., Camilli, G., Maxson, M., Hube, B., Moyes, D. L., Richardson, J. P., and Naglik, J. R. (2021) Candidalysin triggers epithelial cellular stresses that induce necrotic death. Cell Microbiol. 10.1111/CMI.13371

11. Nikou, S. A., Zhou, C., Griffiths, J. S., Kotowicz, N. K., Coleman, B. M., Green, M. J., Moyes, D. L., Gaffen, S. L., Naglik, J. R., and Parker, P. J. (2022) The Candida albicans toxin candidalysin mediates distinct epithelial inflammatory responses through p38 and EGFR-ERK pathways. Sci Signal. 10.11126/SCISIGNAL.ABJ6915

12. Ho, J., Moyes, D. L., Tavassoli, M., and Naglik, J. R. (2017) The Role of ErbB Receptors in Infection. Trends in Microbiology. 25, 942–952

13. Ho, J., Yang, X., Nikou, S. A., Kichik, N., Donkin, A., Ponde, N. O., Richardson, J. P., Gratacap, R. L., Archambault, L. S., Zwirner, C. P., Murciano, C., Henley-Smith, R., Thavaraj, S., Tynan, C. J., Gaffen, S. L., Hube, B., Wheeler, R. T., Moyes, D. L., and Naglik, J. R. (2019) Candidalysin activates innate epithelial immune responses via epidermal growth factor receptor. Nature Communications. 10, 2297

14. Swidergall, M., Solis, N. v., Millet, N., Huang, M. Y., Lin, J., Phan, Q. T., Lazarus, M. D., Wang, Z., Yeaman, M. R., Mitchell, A. P., and Filler, S. G. (2021) Activation of EphA2-EGFR signaling in oral epithelial cells by Candida albicans virulence factors. PLoS Pathogens. 10.1371/journal.ppat.1009221

15. Solis, N. v., Swidergall, M., Bruno, V. M., Gaffen, S. L., and Filler, S. G. (2017) The aryl hydrocarbon receptor governs epithelial cell invasion during oropharyngeal candidiasis. mBio. 8, e00025–17

16. Zhu, W., Phan, Q. T., Boontheung, P., Solis, N. v., Loo, J. A., and Fillera, S. G. (2012) EGFR and HER2 receptor kinase signaling mediate epithelial cell invasion by Candida albicans during oropharyngeal infection. Proc Natl Acad Sci U S A. 109, 14194–14199

17. Ho, J., Wickramasinghe, D. N., Nikou, S.-A., Hube, B., Richardson, J. P., and Naglik, J. R. (2020) Candidalysin Is a Potent Trigger of Alarmin and Antimicrobial Peptide Release in Epithelial Cells. Cells. 9, 699

18. Wee, P., and Wang, Z. (2017) Epidermal growth factor receptor cell proliferation signaling pathways. Cancers (Basel). 9, 52

19. Sigismund, S., Avanzato, D., and Lanzetti, L. (2018) Emerging functions of the EGFR in cancer. Molecular Oncology. 12, 3

20. Shin, D. M., Ro, J. Y., Hong, W. K., and Hittelman, W. N. (1994) Dysregulation of Epidermal Growth Factor Receptor Expression in Premalignant Lesions during Head and Neck Tumorigenesis. Cancer Research

21. Mimuro, H., Suzuki, T., Tanaka, J., Asahi, M., Haas, R., and Sasakawa, C. (2002) Grb2 is a key mediator of Helicobacter pylori CagA protein activities. Molecular Cell. 10, 745–755

22. Mölleken, K., Becker, E., and Hegemann, J. H. (2013) The Chlamydia pneumoniae Invasin Protein Pmp21 Recruits the EGF Receptor for Host Cell Entry. PLoS Pathogens. 9, e1003325

23. Rajaram, M. V. S., Arnett, E., Azad, A. K., Guirado, E., Ni, B., Gerberick, A. D., He, L. Z., Keler, T., Thomas, L. J., Lafuse, W. P., and Schlesinger, L. S. (2017) M. tuberculosis-Initiated Human Mannose Receptor Signaling Regulates Macrophage Recognition and Vesicle Trafficking by FcRγ-Chain, Grb2, and SHP-1. Cell Reports. 21, 126–140

24. Nikou, S. A., Kichik, N., Brown, R., Ponde, N. O., Ho, J., Naglik, J. R., and Richardson, J. P. (2019) Candida albicans interactions with mucosal surfaces during health and disease. Pathogens. 10.3390/pathogens8020053

25. Moyes, D. L., Runglall, M., Murciano, C., Shen, C., Nayar, D., Thavaraj, S., Kohli, A., Islam, A., Mora-Montes, H., Challacombe, S. J., and Naglik, J. R. (2010) A biphasic innate immune MAPK response discriminates between the yeast and hyphal forms of candida albicans in epithelial cells. Cell Host and Microbe. 8, 225–235

26. Moyes, D. L., Shen, C., Murciano, C., Runglall, M., Richardson, J. P., Arno, M., Aldecoa-Otalora, E., and Naglik, J. R. (2014) Protection against epithelial damage during Candida albicans infection is mediated by PI3K/Akt and mammalian target of rapamycin signaling. Journal of Infectious Diseases. 209, 1816–1826

27. Mattoon, D. R., Lamothe, B., Lax, I., and Schlessinger, J. (2004) The docking protein Gab1 is the primary mediator of EGF-stimulated activation of the PI-3K/Akt cell survival pathway. BMC Biology. 2, 24

28. Verma, A. H., Richardson, J. P., Zhou, C., Coleman, B. M., Moyes, D. L., Ho, J., Huppler, R., Ramani, K., McGeachy, M. J., Mufazalov, I. A., Waisman, A., Kane, L. P., Biswas, P. S., Hube, B., Naglik, J. R., and Gaffen, S. L. (2017) Oral epithelial cells orchestrate innate type 17 responses to Candida albicans through the virulence factor candidalysin. Science Immunology. 10.1126/sciimmunol.aam8834

29. Verma, A. H., Zafar, H., Ponde, N. O., Hepworth, O. W., Sihra, D., Aggor, F. E. Y., Ainscough, J. S., Ho, J., Richardson, J. P., Coleman, B. M., Hube, B., Stacey, M., McGeachy, M. J., Naglik, J. R., Gaffen, S. L., and Moyes, D. L. (2018) IL-36 and IL-1/IL-17 Drive Immunity to Oral Candidiasis via Parallel Mechanisms. The Journal of Immunology. 201, 627–634

30. Westman, J., Plumb, J., Licht, A., Yang, M., Allert, S., Naglik, J. R., Hube, B., Grinstein, S., and Maxson, M. E. (2022) Calcium-dependent ESCRT recruitment and lysosome exocytosis maintain epithelial integrity during Candida albicans invasion. Cell Rep. 10.1016/J.CELREP.2021.110187

31. Wu, Y., Zeng, Z., Guo, Y., Song, L., Weatherhead, J. E., Huang, X., Zeng, Y., Bimler, L., Chang, C. Y., Knight, J. M., Valladolid, C., Sun, H., Cruz, M. A., Hube, B., Naglik, J. R., Luong, A. U., Kheradmand, F., and Corry, D. B. (2021) Candida albicans elicits protective allergic responses via platelet mediated T helper 2 and T helper 17 cell polarization. Immunity. 54, 2595–2610.e7

32. Swidergall, M., Khalaji, M., Solis, N. v., Moyes, D. L., Drummond, R. A., Hube, B., Lionakis, M. S., Murdoch, C., Filler, S. G., and Naglik, J. R. (2019) Candidalysin Is Required for Neutrophil Recruitment and Virulence During Systemic Candida albicans Infection. J Infect Dis. 220, 1477–1488

33. Drummond, R. A., Swamydas, M., Oikonomou, V., Zhai, B., Dambuza, I. M., Schaefer, B. C., Bohrer, A. C., Mayer-Barber, K. D., Lira, S. A., Iwakura, Y., Filler, S. G., Brown, G. D., Hube, B., Naglik, J. R., Hohl, T. M., and Lionakis, M. S. (2019) CARD9 + microglia promote antifungal immunity via IL-1β- and CXCL1-mediated neutrophil recruitment. Nat Immunol. 20, 559–570

34. Lemmon, M. A., and Schlessinger, J. (2010) Cell signaling by receptor tyrosine kinases. Cell. 141, 1117–1134

35. Swidergall, M., Solis, N. v., Lionakis, M. S., and Filler, S. G. (2018) EphA2 is an epithelial cell pattern recognition receptor for fungal β-glucans. Nature Microbiology. 3, 53–61

36. Phan, Q. T., Lin, J., Solis, N. v., Eng, M., Swidergall, M., Wang, F., Li, S., Gaffen, S. L., Chou, T. F., and Filler, S. G. (2021) The Globular C1q Receptor Is Required for Epidermal Growth Factor Receptor Signaling during Candida albicans Infection. mBio. 10.1128/MBIO.02716-21

37. Sun, H., Shen, Y., Dokainish, H., Holgado-Madruga, M., Wong, A., and Ireton, K. (2005) Host adaptor proteins Gab1 and CrkII promote InIB-dependent entry of Listeria monocytogenes. Cellular Microbiology. 7, 443–457

38. Batzer, A. G., Rotin, D., Ureña, J. M., Skolnik, E. Y., and Schlessinger, J. (1994) Hierarchy of binding sites for Grb2 and Shc on the epidermal growth factor receptor. Molecular and Cellular Biology. 14, 5192–5201

39. Holgado-Madruga, M., Emlet, D. R., Moscatello, D. K., Godwin, A. K., and Wong, A. J. (1996) A Grb2-associated docking protein in EGF- and insulin-receptor signalling. Nature. 379, 560–564

40. Xiang, Y. P., Xiao, T., Li, Q. G., Lu, S. S., Zhu, W., Liu, Y. Y., Qiu, J. Y., Song, Z. H., Huang, W., Yi, H., Tang, Y. Y., and Xiao, Z. Q. (2020) Y772 phosphorylation of EphA2 is responsible for EphA2-dependent NPC nasopharyngeal carcinoma growth by Shp2/Erk-1/2 signaling pathway. Cell Death and Disease. 10.1038/s41419-020-02831-0

41. Murciano, C., Moyes, D. L., Runglall, M., Tobouti, P., Islam, A., Hoyer, L. L., and Naglik, J. R. (2012) Evaluation of the role of Candida albicans agglutinin-like sequence (ALS) proteins in human oral epithelial cell interactions. PLoS ONE. 10.1371/journal.pone.0033362

42. Prenzel, N., Zwick, E., Daub, H., Leserer, M., Abraham, R., Wallasch, C., and Ullrich, A. (1999) EGF receptor transactivation by G-protein-coupled receptors requires metalloproteinase cleavage of proHB-EGF. Nature. 402, 884–888

43. Zhang, Q., Adiseshaiah, P., and Reddy, S. P. (2005) Matrix metalloproteinase/epidermal growth factor receptor/mitogen-activated protein kinase signaling regulate fra-1 induction by cigarette smoke in lung epithelial cells. American Journal of Respiratory Cell and Molecular Biology. 32, 72–81

44. Keates, S., Sougioultzis, S., Keates, A. C., Zhao, D., Peek, R. M., Shaw, L. M., and Kelly, C. P. (2001) cag+ Helicobacter pylori Induce Transactivation of the Epidermal Growth Factor Receptor in AGS Gastric Epithelial Cells. Journal of Biological Chemistry. 276, 48127–48134

45. Na, X., Zhao, D., Koon, H. W., Kim, H., Husmark, J., Moyer, M. P., Pothoulakis, C., and Lamont, J. T. (2005) Clostridium difficile toxin B activates the EGF receptor and the ERK/MAP kinase pathway in human colonocytes. Gastroenterology. 128, 1002–1011

46. Taylor, J. L., Hattle, J. M., Dreitz, S. A., Troudt, J. L. M., Izzo, L. S., Basaraba, R. J., Orme, M., Matrisian, L. M., and Izzo, A. A. (2006) Role for matrix metalloproteinase 9 in granuloma formation during pulmonary Mycobacterium tuberculosis infection. Infect Immun. 74, 6135–6144

47. Nishikaku, A. S., Ribeiro, L. C., Molina, R. F. S., Albe, B. P., Cunha, C. D. S., and Burger, E. (2009) Matrix metalloproteinases with gelatinolytic activity induced by Paracoccidioides brasiliensis infection. International Journal of Experimental Pathology. 90, 527

48. Bouillot, S., Reboud, E., and Huber, P. (2018) Functional consequences of calcium influx promoted by bacterial pore-forming Toxins. Toxins (Basel). 10.3390/toxins10100387

49. Hsuan, S. L., Kannan, M. S., Jeyaseelan, S., Prakash, Y. S., Sieck, G. C., and Maheswaran, S. K. (1998) Pasteurella haemolytica A1-derived leukotoxin and endotoxin induce intracellular calcium elevation in bovine alveolar macrophages by different signaling pathways. Infection and Immunity. 66, 2836–2844

50. Laestadius, Å., Richter-Dahlfors, A., and Aperia, A. (2002) Dual effects of Escherichia coli α-hemolysin on rat renal proximal tubule cells. Kidney International. 62, 2035–2042

51. Migliaccio, E., Giogio, M., Mele, S., Pelicci, G., Reboldi, P., Pandolfi, P. P., Lanfrancone, L., and Pelicci, P. G. (1999) The p66(shc) adaptor protein controls oxidative stress response and life span in mammals. Nature. 402, 309–313

52. Wang, H. C., Zhou, Y., and Huang, S. K. (2017) SHP-2 phosphatase controls aryl hydrocarbon receptor-mediated ER stress response in mast cells. Archives of Toxicology. 91, 1739–1748

53. Takahashi-Tezuka, M., Yoshida, Y., Fukada, T., Ohtani, T., Yamanaka, Y., Nishida, K., Nakajima, K., Hibi, M., and Hirano, T. (1998) Gab1 Acts as an Adapter Molecule Linking the Cytokine Receptor gp130 to ERK Mitogen-Activated Protein Kinase. Molecular and Cellular Biology. 18, 4109–4117

54. Ward, A. C., Monkhouse, J. L., Hamilton, J. A., and Csar, X. F. (1998) Direct binding of Shc, Grb2, SHP-2 and p40 to the murine granulocyte colony-stimulating factor receptor. Biochimica et Biophysica Acta - Molecular Cell Research. 1448, 70–76

55. Whitney, P. G., Bär, E., Osorio, F., Rogers, N. C., Schraml, B. U., Deddouche, S., LeibundGut-Landmann, S., and Reis e Sousa, C. (2014) Syk signaling in dendritic cells orchestrates innate resistance to systemic fungal infection. PLoS Pathog. 10.1371/JOURNAL.PPAT.1004276

56. Basu, S., Quilici, C., Zhang, H. H., Grail, D., and Dunn, A. (2009) Mice lacking both G-CSF and IL-6 are more susceptible to Candida albicans infection: Critical role of neutrophils in defense against Candida albicans. http://dx.doi.org/10.1080/08977190801987513. 26, p23–34

57. Ouyang, W., Liu, C., Pan, Y., Han, Y., Yang, L., Xia, J., and Xu, F. (2020) SHP2 deficiency promotes Staphylococcus aureus pneumonia following influenza infection. Cell Prolif. 10.1111/CPR.12721

58. Zhao, L., Xia, J., Li, T., Zhou, H., Ouyang, W., Hong, Z., Ke, Y., Qian, J., and Xu, F. (2016) Shp2 Deficiency Impairs the Inflammatory Response Against Haemophilus influenzae by Regulating Macrophage Polarization. The Journal of Infectious Diseases. 214, 625–633

59. An, H., Zhao, W., Hou, J., Zhang, Y., Xie, Y., Zheng, Y., Xu, H., Qian, C., Zhou, J., Yu, Y., Liu, S., Feng, G., and Cao, X. (2006) SHP-2 Phosphatase Negatively Regulates the TRIF Adaptor Protein-Dependent Type I Interferon and Proinflammatory Cytokine Production. Immunity. 25, 919–928

60. de Melker, A. A., van der Horst, G., Calafat, J., Jansen, H., and Borst, J. (2001) C-Cbl ubiquitinates the EGF receptor at the plasma membrane and remains receptor associated throughout the endocytic route. Journal of Cell Science. 114, 2167–2178

61. Grøvdal, L. M., Stang, E., Sorkin, A., and Madshus, I. H. (2004) Direct interaction of Cbl with pTyr 1045 of the EGF receptor (EGFR) is required to sort the EGFR to lysosomes for degradation. Experimental Cell Research. 300, 388–395

62. Sigismund, S., Algisi, V., Nappo, G., Conte, A., Pascolutti, R., Cuomo, A., Bonaldi, T., Argenzio, E., Verhoef, L. G. G. C., Maspero, E., Bianchi, F., Capuani, F., Ciliberto, A., Polo, S., and di Fiore, P. P. (2013) Threshold-controlled ubiquitination of the EGFR directs receptor fate. EMBO Journal. 32, 2140–2157

63. Perez Verdaguer, M., Zhang, T., Paulo, J. A., Gygi, S., Watkins, S. C., Sakurai, H., and Sorkin, A. (2021) Mechanism of p38 MAPK-induced EGFR endocytosis and its crosstalk with ligand-induced pathways. 10.1083/jcb.202102005

64. Rupniak, H. T., Rowlatt, C., Lane, E. B., Steele, J. G., Trejdosiewicz, L. K., Hill, B. T., Laskiewicz, B., and Povey, S. (1985) Characteristics of four new human cell lines derived from squamous cell carcinomas of the head and neck. J Natl Cancer Inst. 75, 621–635

