## Supplementary Figures for "EGFR-MAPK adaptor proteins mediate the epithelial response to *Candida albicans* via the cytolytic peptide toxin, candidalysin"

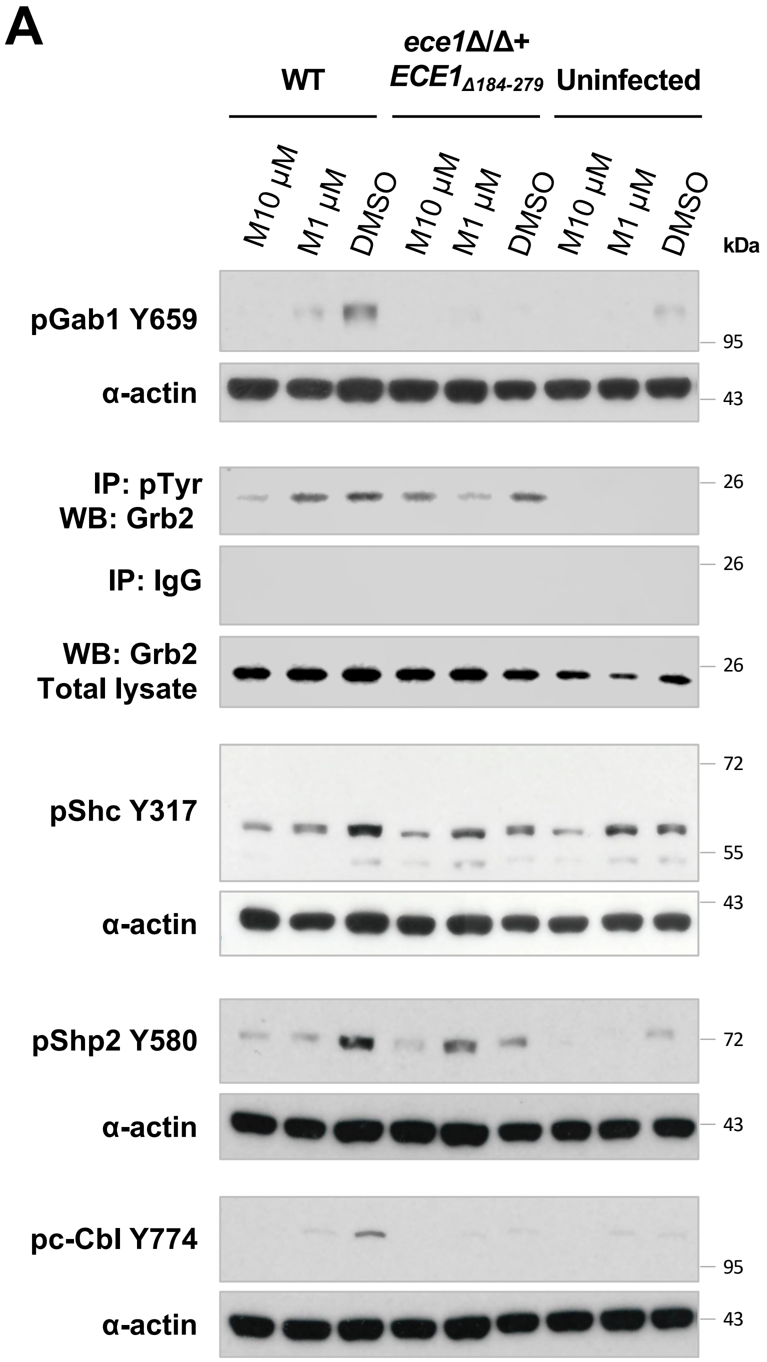


**Figure S1 MMPs are required for *C. albicans*-induced adaptor activation.** MMP inhibition suppresses *C. albicans*-induced adaptor activity. Pre-treatment of TR146 cells with Marimastat (MMP inhibitor, 10 or 1 µM (M10 or M1)) significantly decreased phosphorylation of Gab1, Grb2, Shc, Shp2 and c-Cbl following *C. albicans* infection. Immunoblots are representative of 2 (Grb2) or 3 (Gab1, Shc, Shp2 and c-Cbl) independent experiments.

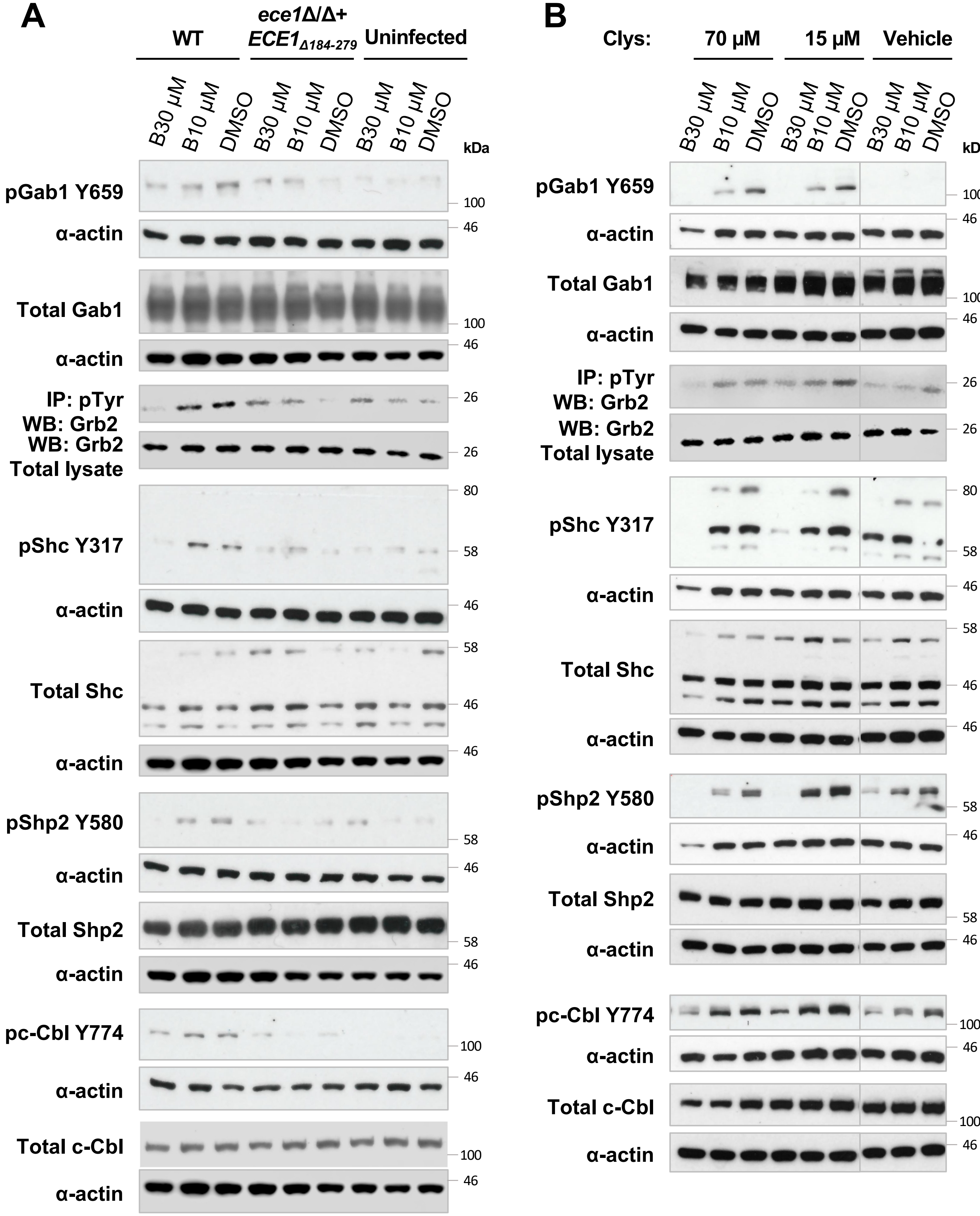


**Figure S2 Candidalysin-induced adaptor activation is driven by calcium flux.** Pre-treatment of TR146 cells with a calcium chelator (BAPTA-AM 30 or 10 μM (B30 or B10)) reduced phosphorylation of Gab1, Grb2, Shc, Shp2 and c-Cbl following A) *C. albicans* infection or B) candidalysin exposure. Protein lysates were taken 2 h post infection with *C. albicans* or candidalysin stimulation for western blot analysis. Immunoblots are representative of 2 (Grb2) or 3 (Gab1, Shc, Shp2 and c-Cbl) independent experiments.

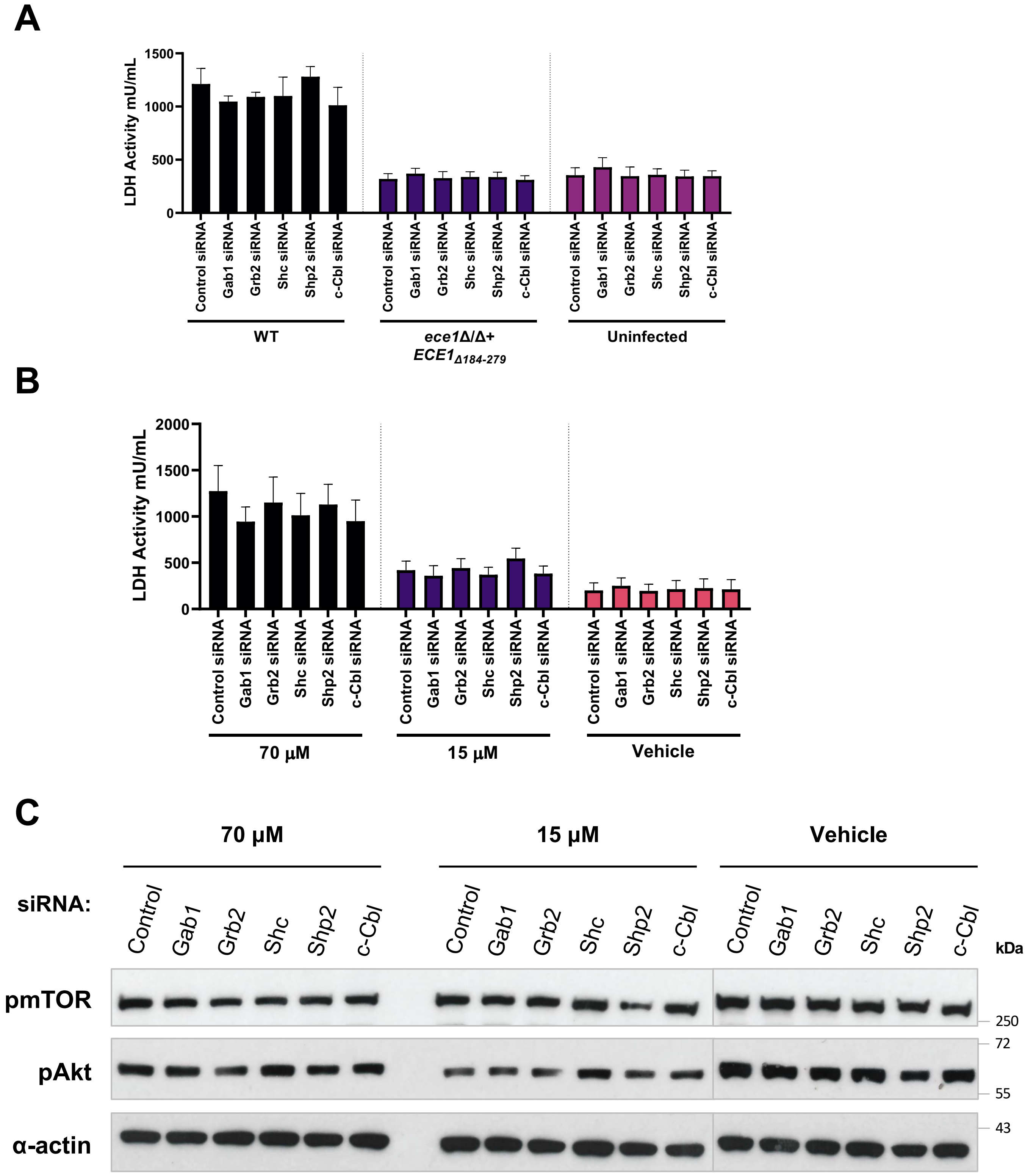


**Figure S3 Adaptors are indispensable in protecting from damage or cell survival in response to candidalysin induced epithelial activation.** Knockdown of Gab1, Grb2, Shc, Shp2 or c-Cbl had no effect on (A-B) LDH release or (C) cell survival pathways. A and B) Following siRNA mediated knockdown, cells were infected with the indicated strains or stimulated with candidalysin and LDH in supernatants evaluated after 24 h. Data are representative of 3 independent experiments. C) TR146 cells were transfected with the indicated siRNAs and stimulated for 2 h with candidalysin (70 or 15 µM) or vehicle. Lysates were immunoblotted for phospho-mTOR (p-mTOR), phospho-Akt (p-Akt) or α-actin. Data are representative of 2 independent experiments. Data were analysed using one way ANOVA with Bonferroni's multiple comparisons test.

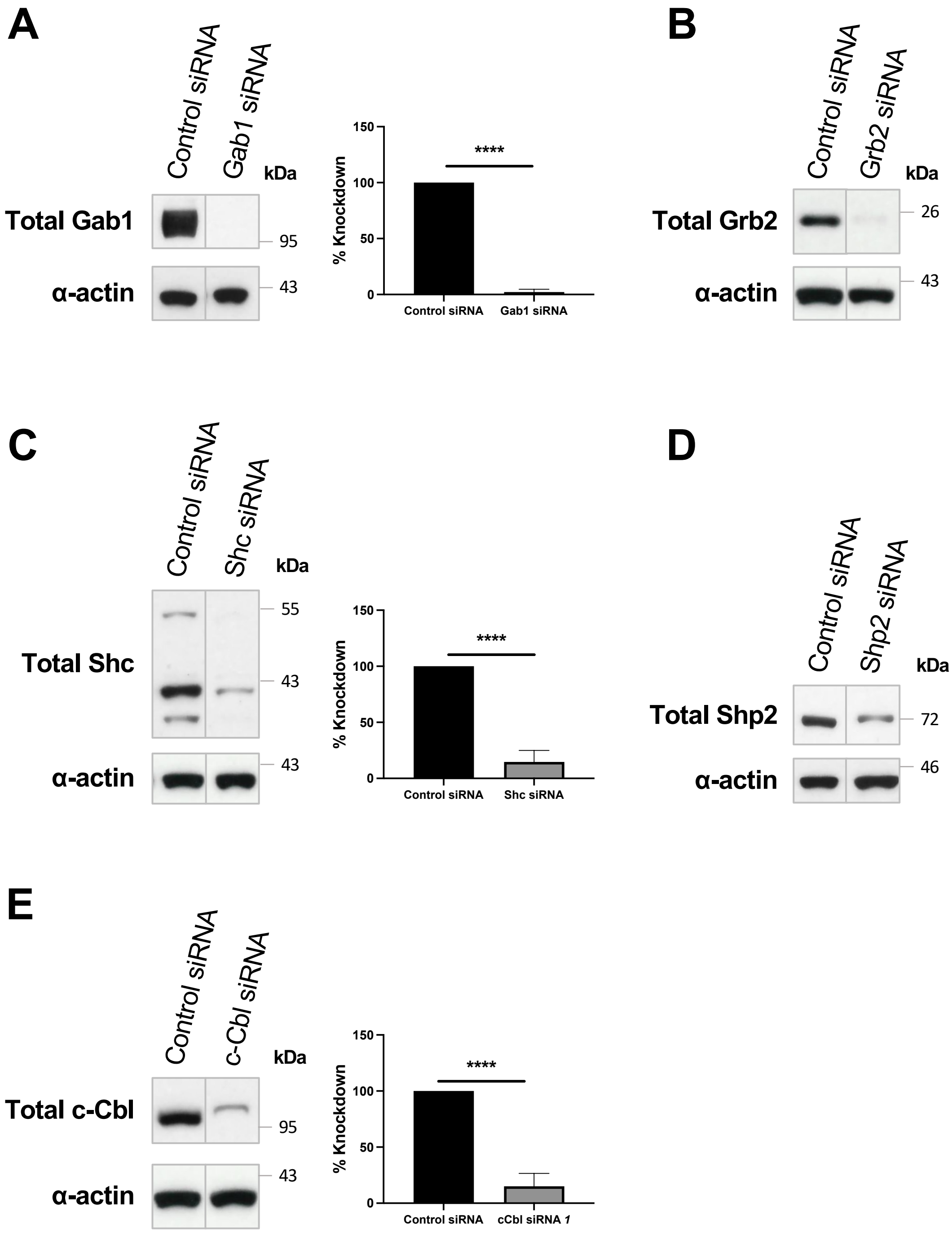


**Figure S4 Optimisation of adaptor siRNA concentration in TR146 oral epithelial cells.** TR146 oral epithelial cells (OECs) were transfected with 5 or 10 nM of the indicated siRNAs for 48 h. Following transfection, cells were lysed and equal amounts of lysate were subjected to SDS-PAGE and western blot analysis with antibodies directed against total (A) Gab1, (B) Grb2, (C) Shc, (D) Shp2 or (E) c-Cbl, as indicated. Data are representative of three biological replicates. Densitometry analysis of westerns was used to assess % knockdown; data are expressed as a percentage of knockdown relative to control siRNA. Data are the mean (±SD) of three biological replicates. Statistical significance was assessed by one way ANOVA with Bonferroni's multiple comparisons test; ** P<0.01, **** P<0.0001.

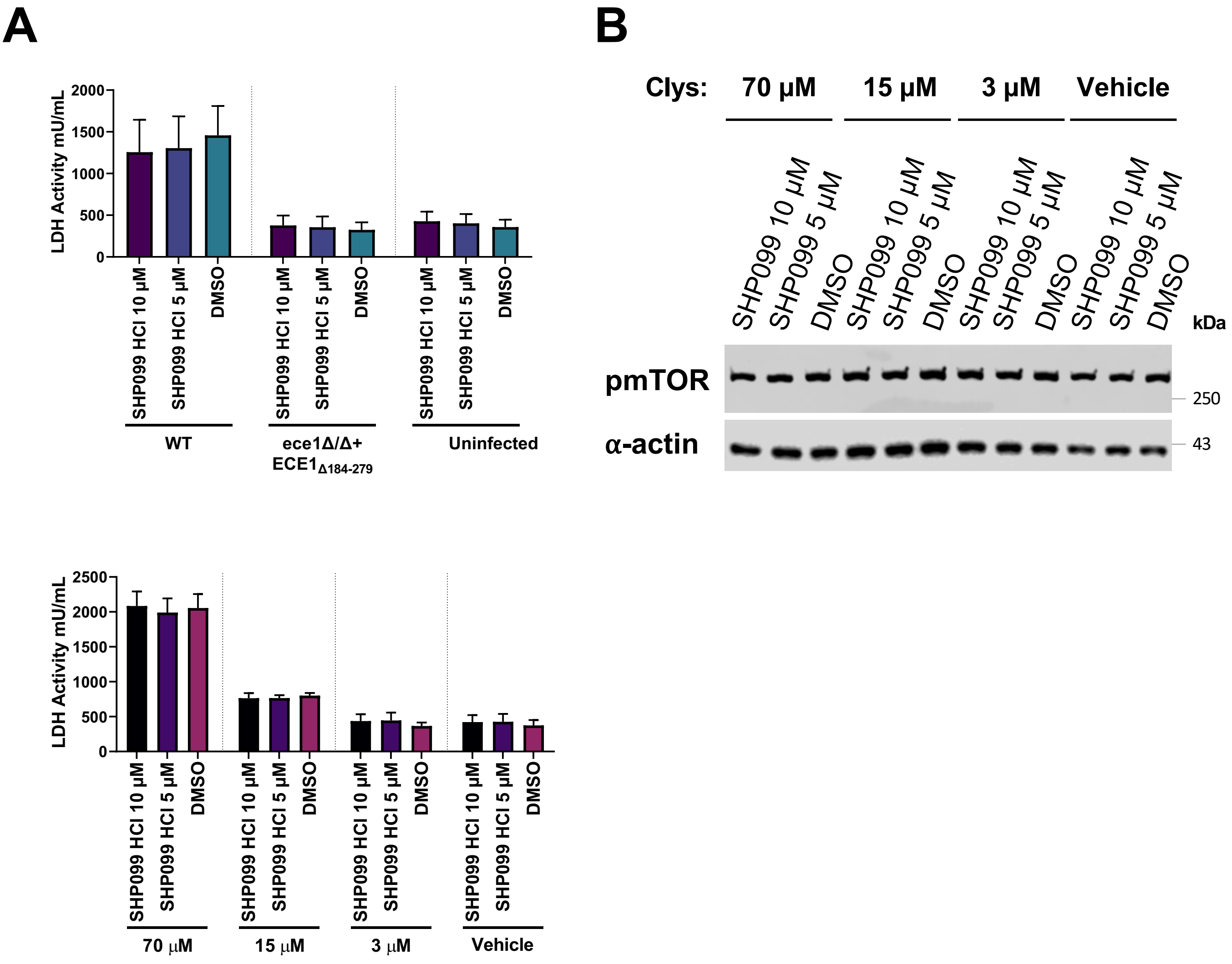


**Figure S5 Shp2 does not protect epithelial cells from damage or activate cell survival pathways in response to candidalysin.** Inhibition of Shp2 activity had no effect on (A) LDH release or (B) cell survival pathways. Data are representative of 3 independent experiments. Data were analysed using one way ANOVA with Bonferroni's multiple comparisons test.

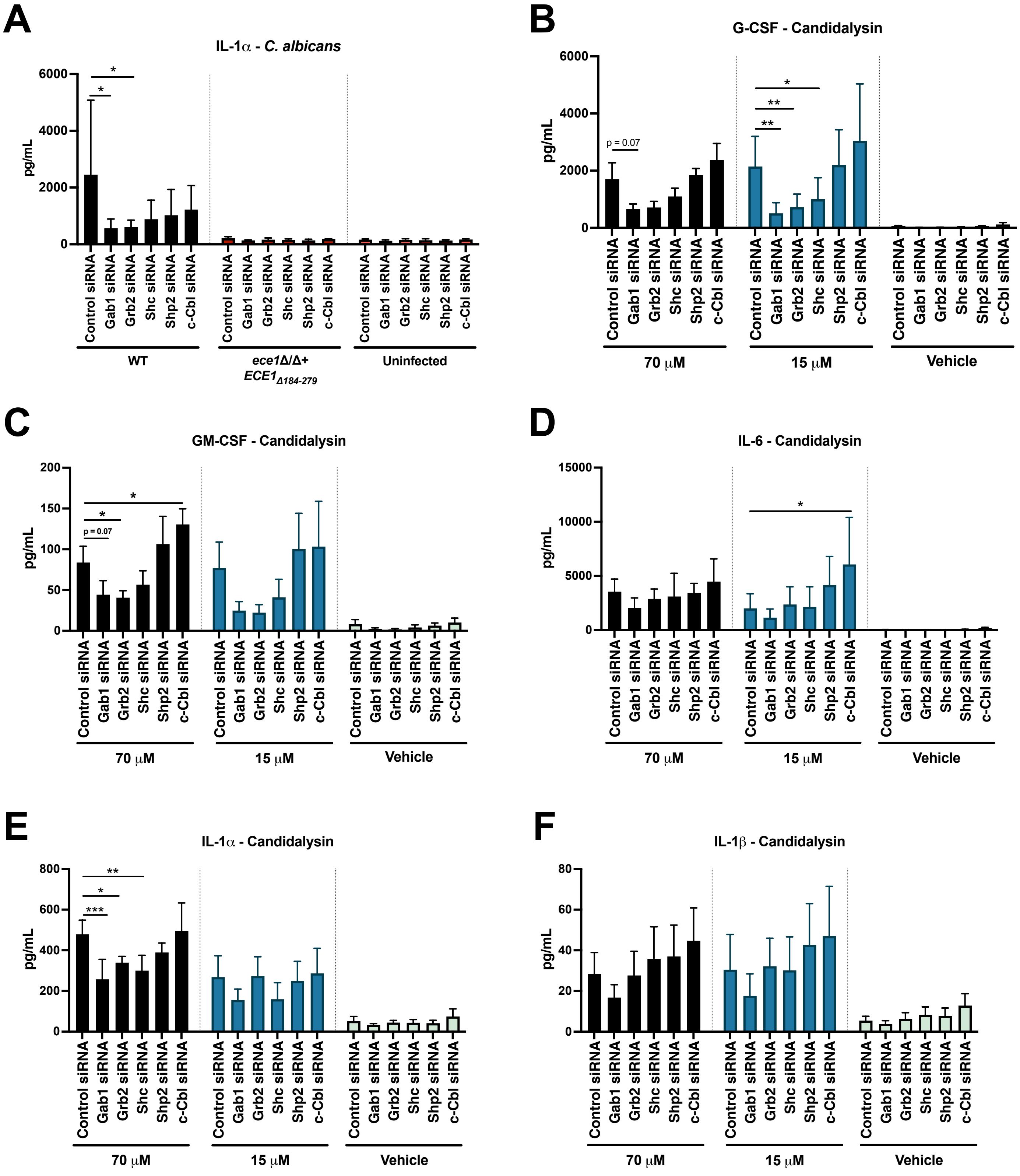


**Figure S6 Adaptors mediate *C. albicans- and* candidalysin-induced cytokine secretion in oral epithelial cells.** Knockdown of Gab1, Grb2 or Shc suppresses G-CSF, GM-CSF, IL-6, IL-1α and IL-1β secretion. A-F) Following siRNA mediated knockdown cells were either infected with the indicated strains or stimulated with candidalysin and supernatants collected after 24 h. Samples were analysed by magnetic Luminex assay to measure cytokine secretion. Data are the mean (±SD) of three biological replicates. Statistical significance was assessed by one way ANOVA with Bonferroni's multiple comparisons test; *P < 0.05, **P < 0.01, ***P<0.001.

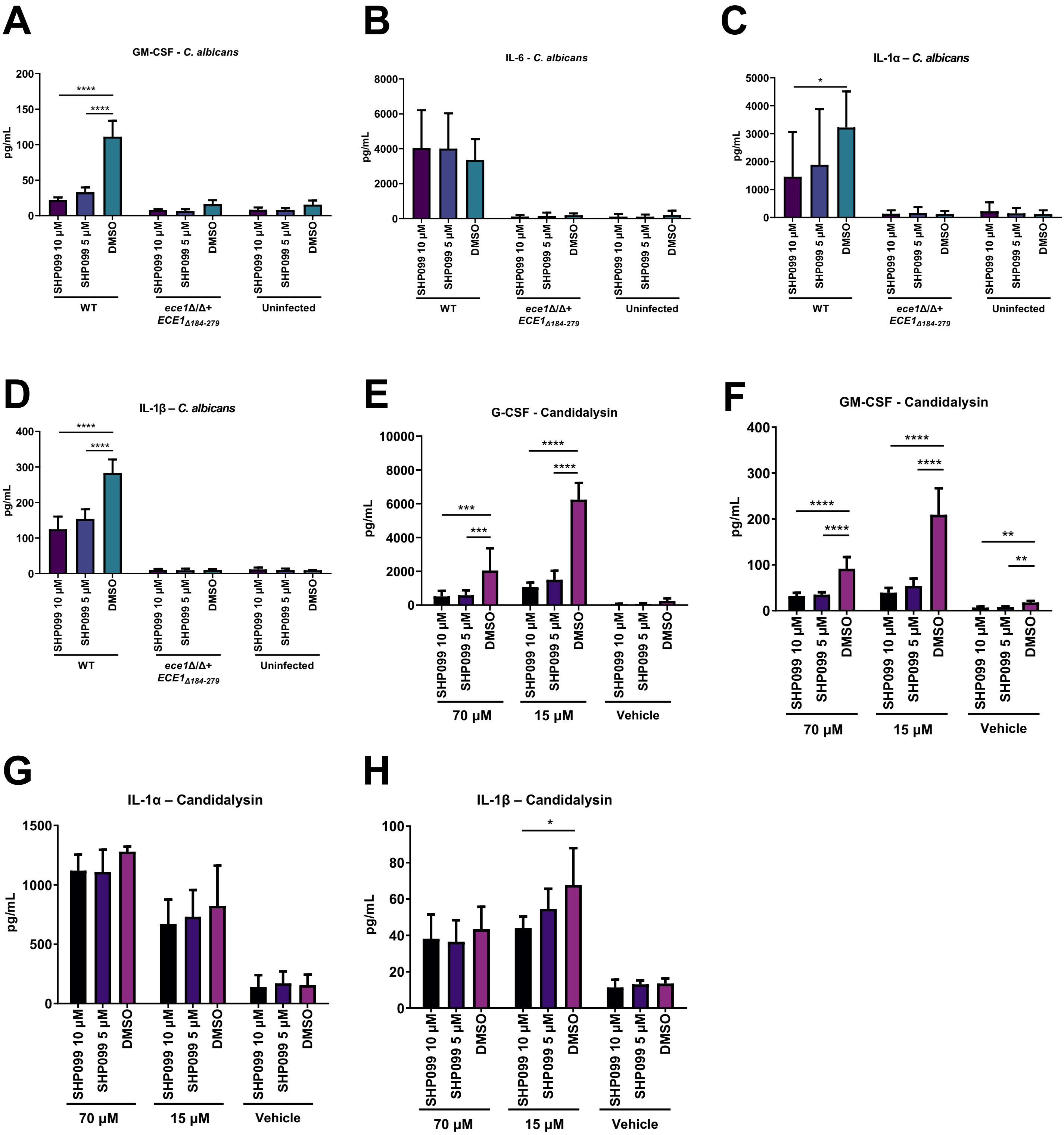


**Figure S7 Adaptors mediate *C. albicans- and* candidalysin-induced cytokine secretion in oral epithelial cells.** Inhibition of Shp2 activity suppresses G-CSF, GM-CSF, IL-1α and IL-1β, but increase IL-6 secretion. A-E) Following pre-treatment with the Shp2 inhibitor, SHP099HCL for 1 h, cells were either infected with the indicated strains or stimulated with candidalysin and supernatants collected after 24 h. Samples were analysed by magnetic Luminex assay to measure cytokine secretion. Data are the mean (±SD) of three biological replicates. Statistical significance was assessed by one way ANOVA with Bonferroni's multiple comparisons test; *P < 0.05, **P < 0.01, ****P<0.0001.

**A**

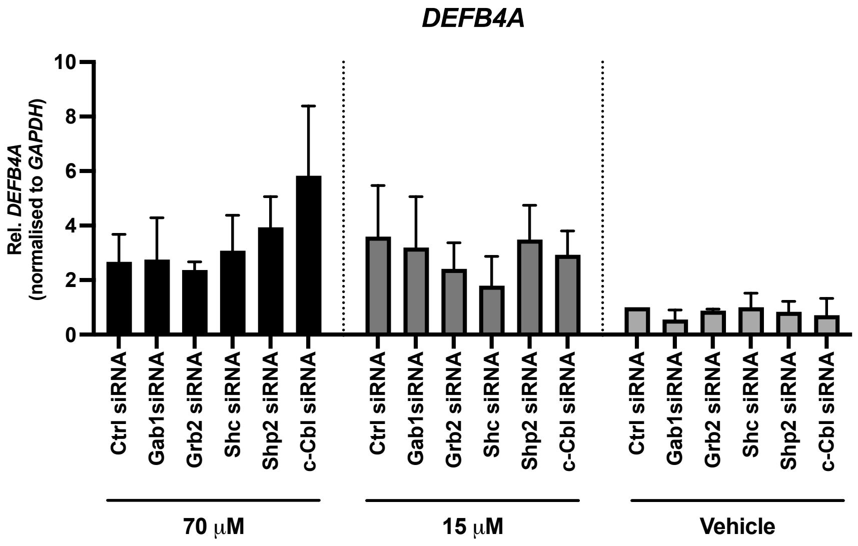


**B**

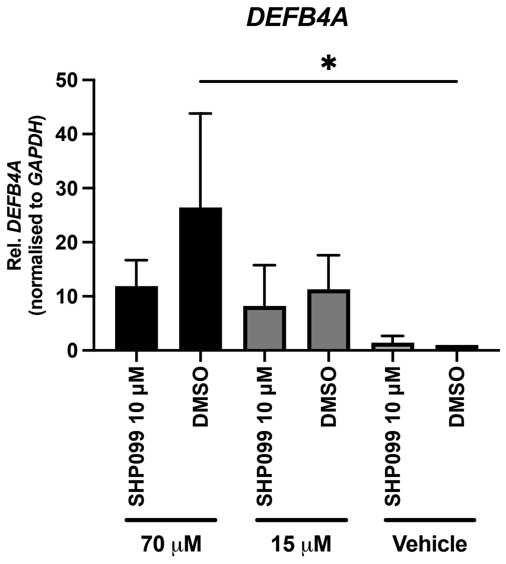


**Figure S8 Adaptors do not mediate *C.* candidalysin-induced *DEFB4A* gene expression in oral epithelial cells.** A) Following siRNA mediated knockdown or B) inhibition of Shp2, cells were stimulated with candidalysin and RNA collected after 6 h. RNA was extracted, cDNA synthesised and gene expression analysed by qPCR. Data are the mean (±SD) of three biological replicates and normalized to control siRNA or DMSO of vehicle. Statistical significance was assessed by one way ANOVA with Bonferroni's multiple comparisons test.

**Table S1: Sequence of siRNAs for knockdown studies**

| siRNA | ID | Sense Sequence | Antisense Sequence |
|---|---|---|---|
| GAB1 | s5463 | CCACGUAAGCAAAAGAGCAtt | UGCUCUUUUGCUUACGUGGtg |
| GRB2 | s226232 | GGUGGAUUAUCACAGAUCUtt | AGAUCUGUGAUAAUCCACCag |
| SHC | s12813 | GCUUUGAAAGUGUCAGUCAtt | UGACUGACACUUUCAAAGCgg |
| PTPN11/SHP2 | s11525 | CAAUGACGGCAAGUCUAAAtt | UUUAGACUUGCCGUCAUUGct |
| CBL | s2476 | GAAUCAACUCUGAACGGAAtt | UUCCGUUCAGAGUUGAUUCtc |
